# G4-iM Grinder: When size and frequency matter. G-Quadruplex, i-Motif and higher order structure search and analysis tool

**DOI:** 10.1101/532382

**Authors:** Efres Belmonte-Reche, Juan Carlos Morales

## Abstract

We present G4-iM Grinder, a system for the localization, characterization and selection of potential G4s, i-Motifs and higher-order structures. A robust and highly adaptable search engine identifies all structures that fit the user’s quadruplex definitions. Their biological relevance, *in-vitro* formation probability and presence of known-to-form structures are then used as filters. The outcome is an efficient methodology that helps select the best candidates for a subsequent *in-vitro* analysis or a macroscopic genomic quadruplex assessment.

As proof of the analytical capabilities of G4-iM Grinder, the human genome was analysed for potential G4s and i-Motifs. Many known-to-form structures were identified. New candidates were selected considering their score and appearance frequency. We also focused on locating Potential Higher Order Quadruplex Sequences (PHOQS). We developed a new methodology to predict the most probable subunits of these assemblies and applied it to a PHOQS candidate.

Taking the human average density as reference, we examined the genomes of several etiological causes of disease. This first of its class comparative study found many organisms to be very dense in these potential quadruplexes. Many presented already known-to-form G4s and i-Motifs. These findings suggest the potential quadruplex have in the fight against these organisms that currently kill millions worldwide.

**GRAPHICAL ABSTRACT:** 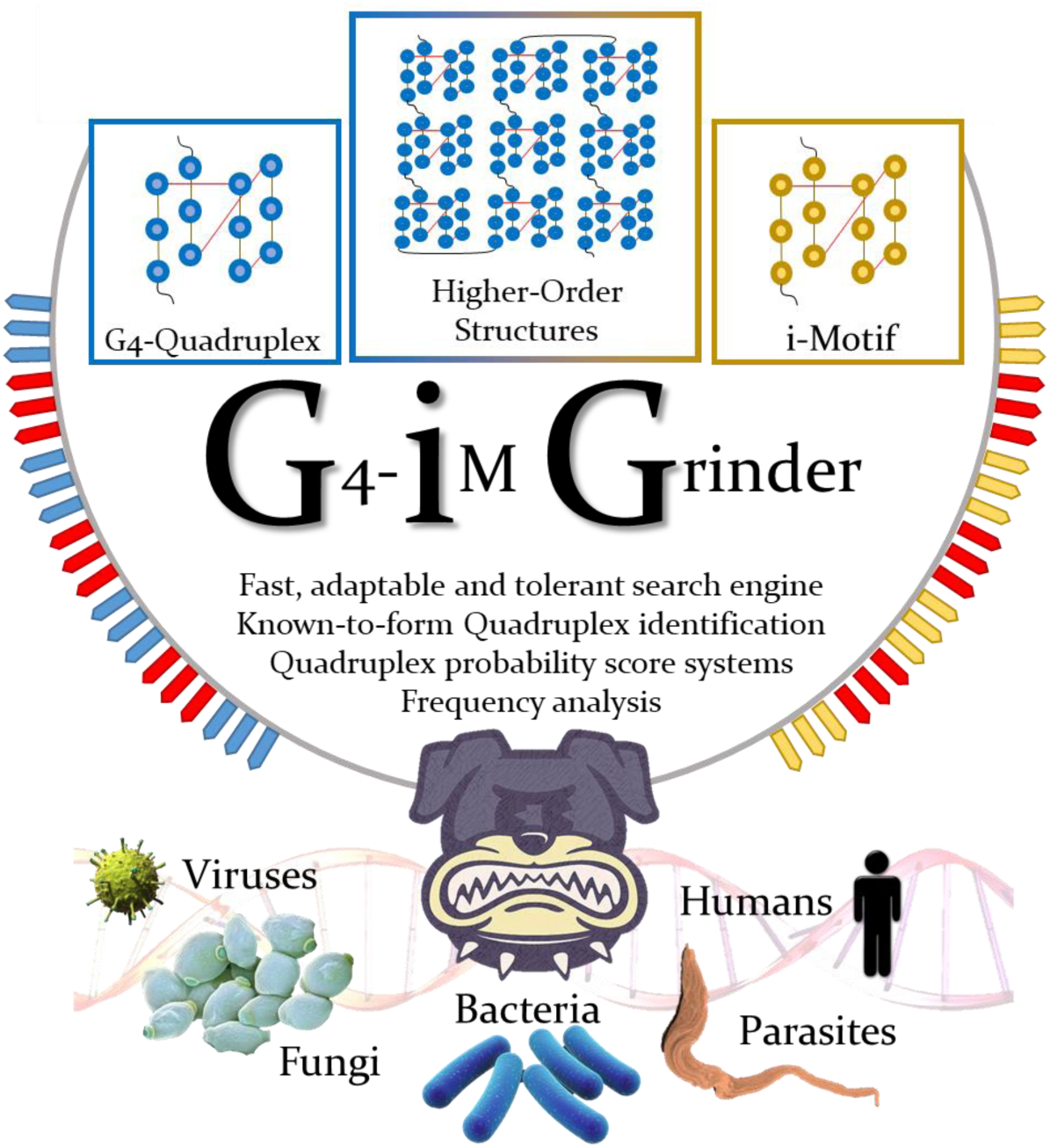

## INTRODUCTION

Guanine rich nucleic acid sequences are capable of forming four-stranded structures called G-quadruplexes (G4), whilst cytosine based assemblies can form i-Motifs. These DNA and RNA conformations have been studied abundantly in the last few years due to the increasing evidence of their functional role in many living organisms (1, 2), yet the natural properties by which they form and work are very much unknown. To identify new structures, *in silico* predictions are based on *in vitro* verified paradigms (3–5). Loops (6), tetrad number, run imperfections (7) and the flanking regions of the structures (5, 8) all seem to play important roles in the topology and dynamics of these secondary structures.

Several tools for the identification of PQSs (putative G4 sequences) within a given DNA/RNA sequence are accessible to users nowadays. The first engines, such as Quadparser (9) and Quadfinder (10), were based on the folding rule which postulates that four perfect G-runs with shorter loops form the most stable G4s. Hence, results with these algorithms yield structures that usually fit the formula: (**G-run** {3:5} Loop {1:7})3 **G-run** {3:5}, where the numbers inside the curly brackets are the range of acceptable lengths of the element.

However, many G4s have been identified that do not follow the folding rule. Loop range unconformity, G-run mismatches and bulges have been confirmed in several G4s (7), so a second generation of PQS search engines was designed to include them in the detection process.

QGRS Mapper (3, 11) partially addressed these irregularities by relaxing the folding rule to accept G-runs of size 2 and loop lengths of up to 45. The likelihood of G4 formation for each result is defined here through a scoring system that favours short and equal loop lengths and higher quartet presence. Similarly, Quadbase2 (12), ImGQfinder (13) and PQSfinder (4) also follow the folding rule (or a similar regular expression model). Of these, Quadbase2 and ImGQfinder are the more basic search engines that heavily restrict user-defined variable configuration. Quadbase2 can detect a fixed number of bulges within the G-runs of predefined size (3) following a regular expression model, and ImGQfinder considers both mismatches and bulges within G-runs in variating G-run sizes. PQSfinder, to the contrary, grants greater parameter liberty and at the same time tolerates G-run defects, such as bulges and mismatches, in the detection process. Its scoring system has been proven to outmatch that of QGRS Mapper and is able to reduce false positive (PQS which are assumed to form G4 but do not) and false negative results (PQS which are assumed to be unable to form G4 but do). PQSfinder is also able to identify and resolve overlapping PQS, which is of utmost importance, as many G4 sequences overlap and compete for the common nucleotides to form the final structures (14).

Search engines that use the sliding window method and break with the folding rule have also been developed and used to detect potential G4s in a genome. Both G4 potential calculator (15) and G4Hunter (16) use this statistical analysis window that willingly defines neither individual PQS boundaries nor defect types. Hence, they can accommodate all G4-*errors* in the search at the expense of being unable to examine overlapping structures (as portions of nucleotides are analysed instead of regular sequences). Results found with G4 potential calculator are then analysed by their G-run density to determine G4-formation potential in a length independent manner. G4Hunter scoring system instead evaluates the result’s G-richness and C-skewness to also consider the experimental destabilization effect caused by nearby cytosine presence on the G-quadruplex (as C can base pair with G and ultimately hinder G-quartet formation (17)).

The newest approach in the field is the development and use of G4-potential scoring methods based on machine-learning algorithms. These avoid predefined motif definitions and minimize formation assumptions to improve the analytical accuracy on non-standard PQSs, at the cost of obscurity in their predictive features. G4NN for example (18), employs an artificial neural network to classify the results of a sliding window model into forming and not-forming RNA G4 sequences. In a similar fashion, Quadron uses an artificial intelligence to classify folding rule abiding PQSs which returns the longest results for all the possible nested G4 sequences (5).

All quadruplex search models have several drawbacks and limitations despite the advances in the field. For the most part, variable configuration is usually heavily restricted meaning only the same kind of structures can be looked for (Table 1), excluding -for example- the detection of structures with more than four G-runs in the sequence. Even if only four G-runs can form the G4-tetrads, extra G-runs can also occur in G-quadruplexes (19, 20) or as part of a fluctuating structure (21). Additionally, no current search engine considers nor calculates genomic PQS frequency from the results. Even if a higher frequency of a PQS does not mean a stronger tendency of *in vitro* G4 formation, it does mean that statistically they may be more biologically important or less biologically problematic. Also, higher G4 frequency allows easier and more accessible targets for the current G4-ligands, which in general are not selective between G4s (22–24). As example, we recently published the results of a PQS search in several parasitic genomes whilst considering frequency, and identified numerous highly recurrent potential G4 candidates (25). Most of these had already been described in literature as G4-forming structures; yet other sequences were new, including EBR1 which is repeated 33 times in the genome of *Trypanosoma brucei.* Despite EBR1 being graded poorly by the engine employed, we confirmed that the recurrent parasitic PQS was able to form G4 in solution even in the absence of cations. Other examples of the use of frequency as a selection filter also exist (26).

**Table 1.** Comparison of some of the search engines and analysers available for use. Structure qualification includes composition analysis and identification of sequences which are already known to form G4 *in vitro* within the results. Abbreviations: F. R. is folding rule, F. F. R. is flexible folding rule, S. W. is sliding window, A. N. N. is artificial neural network. Y stands for Yes, N for No.

|  | **QGRS Mapper** | **G4Hunter** | **PQSfinder** | **G4RNA Screener** | **G4-iM Grinder** |
| --- | --- | --- | --- | --- | --- |
| **Format** | web | R script/web | R package | Python script/web | R package |
| **Search engine** |  |  |  |  |  |
| Model | F.R. | S.W. | F.F.R. | S.W. | F.F.R. |
| Run Composition | G | G, C | G | G | G, C, T, U, A |
| Run Imperfections | N | **Y** | **Y** | **Y** | **Y** |
| Modulable variables | 5 | 2 | 10 | 3 | **13** |
| **Results analysis** |  |  |  |  |  |
| Structure analysis | **Y** | **Y** | **Y** | **Y** | **Y** |
| Structure frequency analysis | N | N | N | N | **Y** |
| High-order search & analysis | N | N | N | N | **Y** |
| Structure Qualification | N | N | N | N | **Y** |
| PQS Score | G-Score | G4Hunter | PQSfinder | G4NN (A. N. N.)<br>G4Hunter<br>cGcC | PQSfinder<br>G4Hunter<br>cGcC<br>Frequency<br>Total Score |

To conclude, none of the search engines have been explicitly designed to detect, analyse and evaluate higher order sequences. These assemblies with great biological potential are the result of very rich genomic G-tracks that form consecutive G4s. These can be assembled into a higher order structure formed by several G4 subunits. The human telomere sequence (hTel) higher order assembly is currently the main focus of this new area of investigation (27–30). Although several different models exist regarding the interactions between the units, the supra-structure has been found to influence the interactions between the hTel G4s and the telomeric proteins compared to individual G4s.

## MATERIAL AND METHODS

### G4-iM Grinder

Our contribution to the field is focused on solving these limitations in an easy and fast manner for the user. G4-iM Grinder is an *in silico* tool designed as the starting step of the genomic quadruplex relevance study workflow. It is intended for researchers who want to detect quadruplex therapeutic targets (both known and new) on a genome, and efficiently filter and select the most interesting results to analyse *in vitro*. The final objective of the algorithm is to save time and resources in finding, evaluating, selecting and confirming these genomic structures.

Three distinct processes constitute G4-iM Grinder: the quadruplex search engine, the quadruplex qualification functions and the quantification functions. These processes tolerate parallelization over several cores to expedite the analysis.

Search engine: G4-iM Grinder’s search engine was developed to allow extensive freedom in the user’s definition of a quadruplex. These parameters are then applied to a fast, reliable and tolerant search motor capable of detecting even potential higher-order structures (Supplementary information – 1. G4-iM Grinder’s search and analyser algorithm). The result is a very flexible algorithm capable of detecting all structures that fit the user’s prerequisites, defined in 13 variables (Supplementary information – 2. Variables, predefined values and examples). These variables all have predefined values that conform to a functional yet broad definition of a flexible folding rule quadruplex disposition (Supplementary information – 3. G-Quadruplex and G4-iM Grinder). They can, however, be easily modified if, for example: structures with longer loops, more run bulges and/or shorter run sizes are also to be detected.

Method 1 (M1), Method 2 (M2) and Method 3 (M3) sub-processes constitute the search engine (Figure 1, A). M1 locates the runs (M1A) and finds their direct run-relationships (M1B), whilst M2 and M3 analyses the potential quadruplex structure formation. M2 does this analysis in an overlapping size-restricted manner (to detect quadruplex structures), and M3 in a <u>non</u>-overlapping size <u>un</u>restricted way (to detect higher order sequences). In each case, the genomic location (M2A and M3A) and the genomic appearance frequency of each sequence is returned (M2B and M3B).

**Figure 1.**
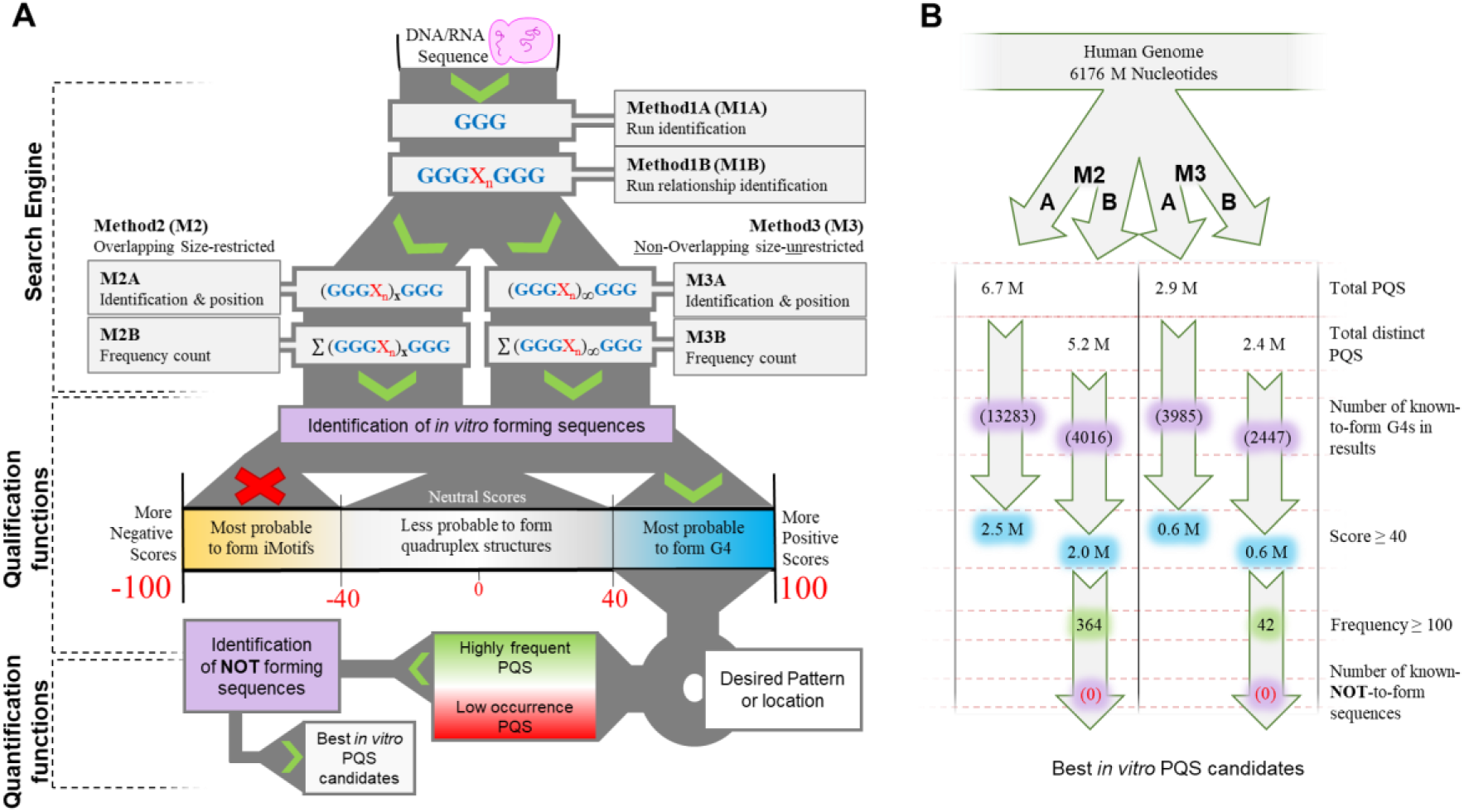
**<u>A</u>**. G4-iM Grinder’s workflow when *RunComposition* = G to find interesting PQS and PHOQS in a genome. Results are filtered by their scores, their frequency and the presence of known-to-form Quadruplex and known-NOT-to-form sequences. **<u>B</u>**. G4-iM Grinder’s workflow results when applied to the human genome. Millions is abbreviated M.

Quadruplex qualification and quantification functions: Several qualification functions were created, updated or adapted to narrow the number of interesting sequences after a search. The traditional qualification approach is to apply scoring systems that link high scores with high *in vitro* probability of formation as a selection or filtering mechanism. G4-iM Grinder incorporates some of these systems (G4hunter and cGcC) to evaluate its results. PQSfinder algorithm was upgraded using machine learning to overcome its current limitations and can also be used for this purpose. It is now capable of evaluating normal as well as irregular quadruplexes, i-Motifs and higher order sequences (Supplementary information – 5. Scoring models and their adaptations). A final score considering a weighted average formula of each scoring system and the frequency of the sequence (modulated by the variables *WeightedParameters* and *FreqWeight* respectively) can also be calculated as the sequence’s quantitative interest score.

Other functions have been integrated into G4-iM Grinder to complement the selection process. The quantification of a predefined pattern as a percentage of the sequence (for example ‘G’, ‘GGG’ or ‘TTA’) and the localization of *in vitro* known-to-form quadruplex and known-**NOT**-to-form sequences can further help select the most interesting sequences (Supplementary information – 6. Explanation of other Variables).

Herein, we propose combining these G4-iM Grinder’s capabilities in an efficient workflow. All sequences found within G4-iM Grinder’s results with known-to-form quadruplex structures are already verified therapeutic targets ready for further investigation. Other sequences should be first filtered by their score. We found that a score of 40 or over is a good threshold that represents over 97 % of all known-to-form G4s examined (Supplementary information – 3. G-Quadruplex and G4-iM Grinder), and therefore represent the most plausible to form sequences. Genomic areas of interest (such as genes) or specific nucleotide arrangements within the quadruplex structure can be further used as filters. Alternatively, a frequency threshold can be employed that redirects the focus to the most recurrent high scoring structures. This way with few biophysical assays, many quadruplex targets with potentially greater biological repercussion can be confirmed. Known-NOT-to-form sequences can then be matched to filter out sequences that coincide. The resulting sequences are therefore the most interesting results to start an quadruplex *in vitro* evaluation.

### i-Motifs and G4-iM Grinder

C-rich regions of DNA or ARN have the ability to fold into tetrameric structures known as i-Motif (31, 32). These DNA assemblies consist of stranded duplexes sustained by hydrogen interactions between the intercalated nucleotide base pairs ***C·C****^+^*, which are stronger than the canonical G-C base pair when under acidic physiological conditions of temperature and ionic strength (33). Several consecutive ***C***s / ***C***^+^s constitutes a run, and four **C-runs** arrange spatially as the final tetramer. The rest of the nucleotides in between **C-runs** form the minor or mayor grooves (loops). As an example, the telomeric sequence (**CCC**ATT)3**CCC** (34), which can be organized as loops and runs, is able to form an i-Motif, yet the rich in C sequence CTCCCTTCTCCTCTC cannot (35).

G4-iM Grinder was designed to allow the search and evaluation of Putative i-Motifs Sequences (PiMS). Its search engine can locate all sequences that can form these quadruplexes using the same flexible folding rule used for G4s, as both are comprised of runs and loops. This follows previous uses of G4 search engines to detect PiMS for *in vitro* evaluation (36).

If the user wishes, G4-iM Grinder can use its qualification functions to evaluate the PiMS’s *in vitro* formation potential. These include the application of the updated *PQSfinder*, *cGcC, G4Hunter* and *Final.Score* scoring algorithms, which operate (for i-Motifs) in an equal but contrary scale to G4s (where bigger <u>negative</u> values mean higher i-Motif probability; bigger <u>positive</u> values mean higher G4 probability).

Potentially, these scoring methods are useful for i-Motif punctuation as the algorithm’s evaluation characteristics (designed for G4s) are also expected to influence C-based structure stability. For example, G4Hunter and cGcC analyse the sequence G and C relationships whilst PQSfinder examines the tetrad size, bulges between tetrads and loop size to assess the potential of the sequences. 95 known-to-form i-Motifs published in literature were located, listed and analysed with G4-iM Grinder to test the use of these scoring systems in i-Motifs (Supplementary information – 4. i-Motifs and G4-iM Grinder). The mean score ± Standard Deviation (SD) of all these i-Motifs were: *G4Hunter* = −49.5 ± 17.0; *PQSfinder* = −62.1 ± 10.7; *score(mean)* = −55.8 ± 13.6, which indicates a strong relationship between the absolute high probability formation scores and the actual *in vitro* formation of the i-Motif (Supplementary information, Figure 4-Top: boxplots). These score relationships are very similar to that of the known-to-form G4s structures (Supplementary information – 3. Score (mean) = 52.9 ± 13.1) that validate the direct link between G4 *in vitro* formation and PQS score. In a similar fashion, these i-Motif results validate the link between i-Motif *in vitro* formation and the PiMS score. Furthermore, 87 % of i-Motifs scored over 40, and given the similar mean and SD to the G4s examined, we propose to use the same threshold of Score ≥ 40 to filter off these results. The pH dependent melting temperatures (Tm) of all available i-Motifs were also listed and used to find potential relationships with their scores and length. An evident direct correlation linking Tm and scores was measured (especially for pH 5, R^2^ = 0.6), despite the deviating experimental conditions of the source’s Tm determination (Supplementary information, Figure 4-bottom: graphs).

## RESULTS

### Full genomic analysis with G4-iM Grinder

An initial performance run was executed on human chromosome 22 to test the performance of the algorithm (Supplementary information – 7. G4-iM Grinder performance). Then, a full analysis of all the human chromosomes was carried out with predefined variable configuration for the identification of G-based PQS and C-based PiMS. Due to the small difference in results between using different *BulgeSize* values (as it is constrained by the variable *MaxIL,* maximum number of bulges in a sequence), it was decided to accept only 1 different nucleotide per run, for a maximum total of 3 per sequence. It is well established that tetrad bulges are a factor for overall structure instability (37) and hence allowing too many of them would result in an increase in the detection rate of low probability *in vitro*-forming structures plus an increase in the computation processing time.

The complete human genomic analysis (M1, M2 and M3, A and B) took 12.6 h for G-based PQS and 10.4 h for C-based PiMS. In both cases, over 6.7 million potential sequences were detected with M2A (overlapping size-restricted method), and 2.9 million with M3A (<u>non</u>-overlapping size-<u>un</u>restricted method - Figure 1, B). These results include thousands of confirmed G-quadruplexes and i-Motifs. Overall, the non-unique sequences (percentage of sequences with a frequency of occurrence of over 1) represent 26 and 21 % of all results (M2A and M3A respectively). Some of these sequences are repeated over 30000 times, although the average is 6.5 repetitions per non-unique sequence.

Following the workflow described previously, results were filtered by their scores to focus on the most probable to form *in vitro* sequences. Doing so for PQS and PiMS reduced the number of results found with M2 threefold whilst maintaining the ratio of unique to non-unique sequences. Of these (over 2 million results), 364 also had a frequency higher than 100, including the confirmed G-Quadruplexes T30695_or_T30923 (38), 20h (39), G4CT-pallidum (40), 93del (41) and 22Ag (42). None of these presented within known-NOT-to-form sequences. Similarly, applying the score filter decreased four fold (to 0.6 million) the original results of M3. Of these 0.6 million, only 42 sequences also had a frequency of at least 100. The most frequent sequences after applying these filters revealed interesting relationships, including the combination of entries 2 and 4 to give the bigger sequences of entries 6 and 7 of Figure 2, B. These combinations identify several widely spread high scoring sequences throughout the human genome, which can potentially form a changeable higher order structure or a fluctuating quadruplex.

**Figure 2.**
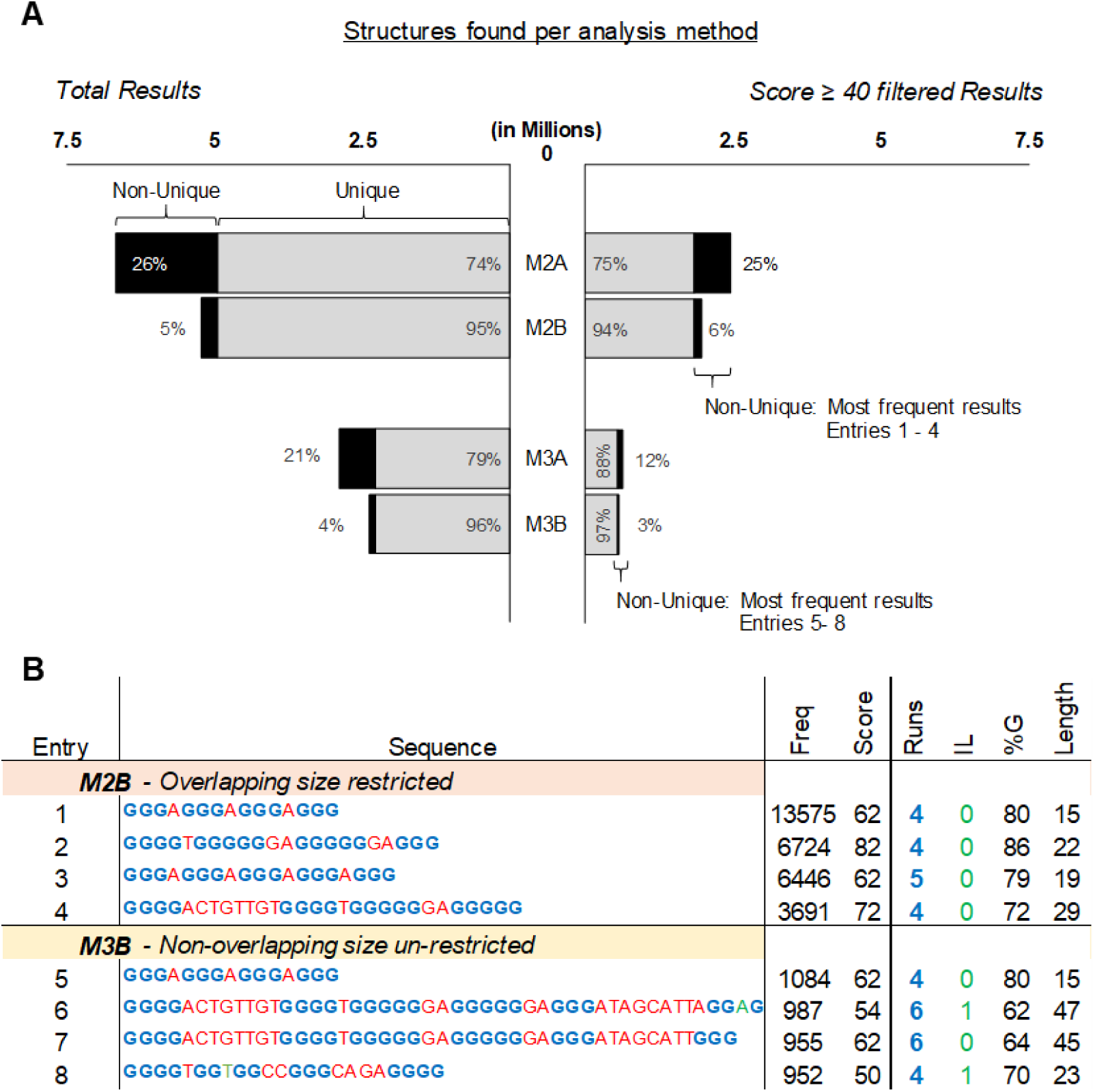
**<u>A.</u>** Total found PQS (left) and found PQS after filtering by score (right) within the entire human genome per method of analysis. The results are divided into unique structures (found with a frequency of 1, in grey), and non-unique structures (found with a frequency of more than 1, in black). The percentage regarding the total is also shown in between parenthesis. **<u>B.</u>** Top four most frequent PQS found within the filtered by score results per method of analysis. Sequence portions in blue are the detected G-runs, in green the run bulges and in red the loops. Score is the average between G4Hunter and PQSfinder. Abbreviations: Freq is frequency, IL is total run bulges and %G is the percentage of G in the sequence.

The search for PiMS gave very similar results to that of PQS. Most noticeably, KRC6 (43), hTel (44), cJun (45) and C3T333 (46) were located within the 357 sequences found with M2 that score at least 40 and have a frequency of at least 100.

### Potential Higher Order Quadruplex Sequences (PHOQS) and their analysis

Using M3A results, the longest of all possible higher order quadruplex sequences (PHOQS) was identified in chromosome 6. This structure potentially involves more than 2700 nucleotides and can be formed by over 300 possible PQS options, yet it was graded poorly because of its many bulges in between G-runs. Hence, the focus was set on the longest structures with the highest probability of formation (Score ≥ 50, Table 2). PHOQS found this way include a 2343 long sequence in chromosome 11 (entry 1) and a 1005 segment in the end of chromosome X, rich in the telomeric and other known-to-form G4 sequences (entry 10).

**Table 2.** Longest Potential Higher Order Quadruplex Sequence (PHOQS) found in the human genome, which scores at least 50 with method 3A (M3A). Conf.Quad.Seqs are identified G4 sequences within the found structure which are known to form *in vitro*, followed by the times detected in between parenthesis. Score is the mean of G4hunter and PQSfinder. Abbreviations: Chrom. is chromosome, IL is run bulges, %G, %C, %A and %T is the percentage of that nucleotide in the sequence.

| Entry | Chrom. | Start | Length | Runs | IL | Strand | Score | %G | %C | %A | %T | Conf. Quad. Seqs |
| --- | --- | --- | --- | --- | --- | --- | --- | --- | --- | --- | --- | --- |
| 1 | 11 | 400747 | 2343 | 305 | 13 | - | 58 | 66 | 17 | 7 | 10 |  |
| 2 | X | 1587610 | 1666 | 293 | 37 | - | 61 | 70 | 1 | 29 | 1 |  |
| 3 | 20 | 64093767 | 1646 | 151 | 13 | + | 53 | 60 | 8 | 13 | 19 |  |
| 4 | 2 | 240923448 | 1351 | 156 | 0 | - | 54 | 59 | 9 | 7 | 25 |  |
| 5 | X | 328170 | 1240 | 182 | 16 | - | 68 | 74 | 5 | 21 | 1 |  |
| 6 | 10 | 131041965 | 1170 | 148 | 2 | + | 56 | 55 | 2 | 22 | 21 |  |
| 7 | 7 | 470172 | 1125 | 136 | 0 | - | 54 | 58 | 8 | 17 | 16 |  |
| 8 | 8 | 141812709 | 1102 | 142 | 8 | - | 54 | 60 | 9 | 13 | 18 |  |
| 9 | 9 | 133668264 | 1089 | 189 | 11 | - | 51 | 66 | 1 | 27 | 6 |  |
| 10 | X | 156029890 | 1005 | 164 | 6 | + | 52 | 56 | 0 | 13 | 31 | 26gtel4 (4) 22Ag (68) Tet22 (7)<br>Gia18 (2) Scer21 (1) 45Ag (51)<br>26gsc (1) |
| 11 | 2 | 239737366 | 990 | 163 | 4 | - | 60 | 63 | 19 | 4 | 14 | d(G4C2)4 (3) |
| 12 | 3 | 10005 | 978 | 159 | 1 | - | 52 | 55 | 2 | 15 | 29 | TSG24 (44) X3ACT (3) Tet22<br>(2) G4CT-pallidum (6) |

Attention was set on HoEBR1, a relatively small sized (< 200 nucleotides to avoid excessive complexity), high scoring and frequent PHOQS. This 118 nucleotide-long PHOQS is repeated four times in the human complementary strand of chromosome 16. Here, it forms part of a nuclear pore complex interacting proteins (NPIPA1 and 2 genes), a polycystin 1 transient receptor potential channel and several other unidentified genes. HoEBR1 can be formed by a combination of its 32 potential PQS subunits (identified by extracting the results from M2A within the location of HoEBR1, Table 3). The known-to-form G4 sequence IV-1242540 was also located within these potential subunits (47).

**Table 3.**
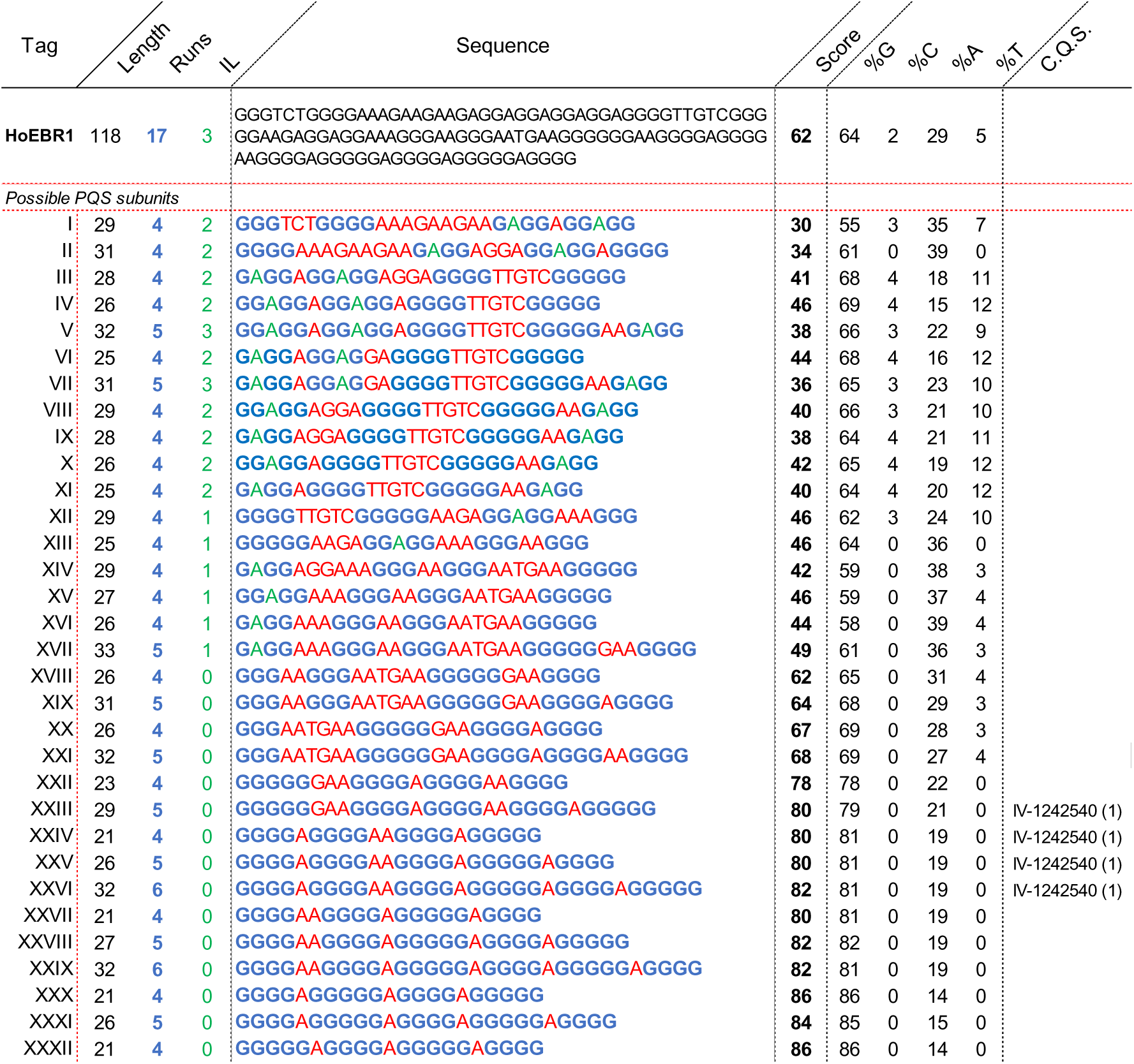
HoEBR1 analysis and dissection into its core possible PQS subunits. In black and in the first row HoEBR1, and beneath are all the possible PQS units that can potentially form the higher order structure. Sequence portions in blue are the detected G-runs, in green the run bulges and in red the loops. Score is the average between G4Hunter and PQSfinder. Abbreviations: C. Q. S. (confirmed quadruplex sequences) are identified G4 structures within the sequence which are known to form *in vitro*, followed by the times detected in between parenthesis, IL is total run bulges, %G, %C, %A and %T is the percentage of that nucleotide in the sequence.

All these subunits overlap and will potentially compete to form the most stable structures. An algorithm was developed to predict the most interesting combinations of PQS subunits to form HoEBR1. Such a tool is included in the G4-iM Grinder package under the function *GiG.M3Structure*. The idea behind the code is to consider the PHOQS as several *seats* for which all the subunits are candidates. When a candidate claims a *seat*, it will annul any other candidate with which it shares nucleotides. In our case, HoEBR1 can be potentially be formed by up to four *seats* (Figure 3, A).

**Figure 3.**
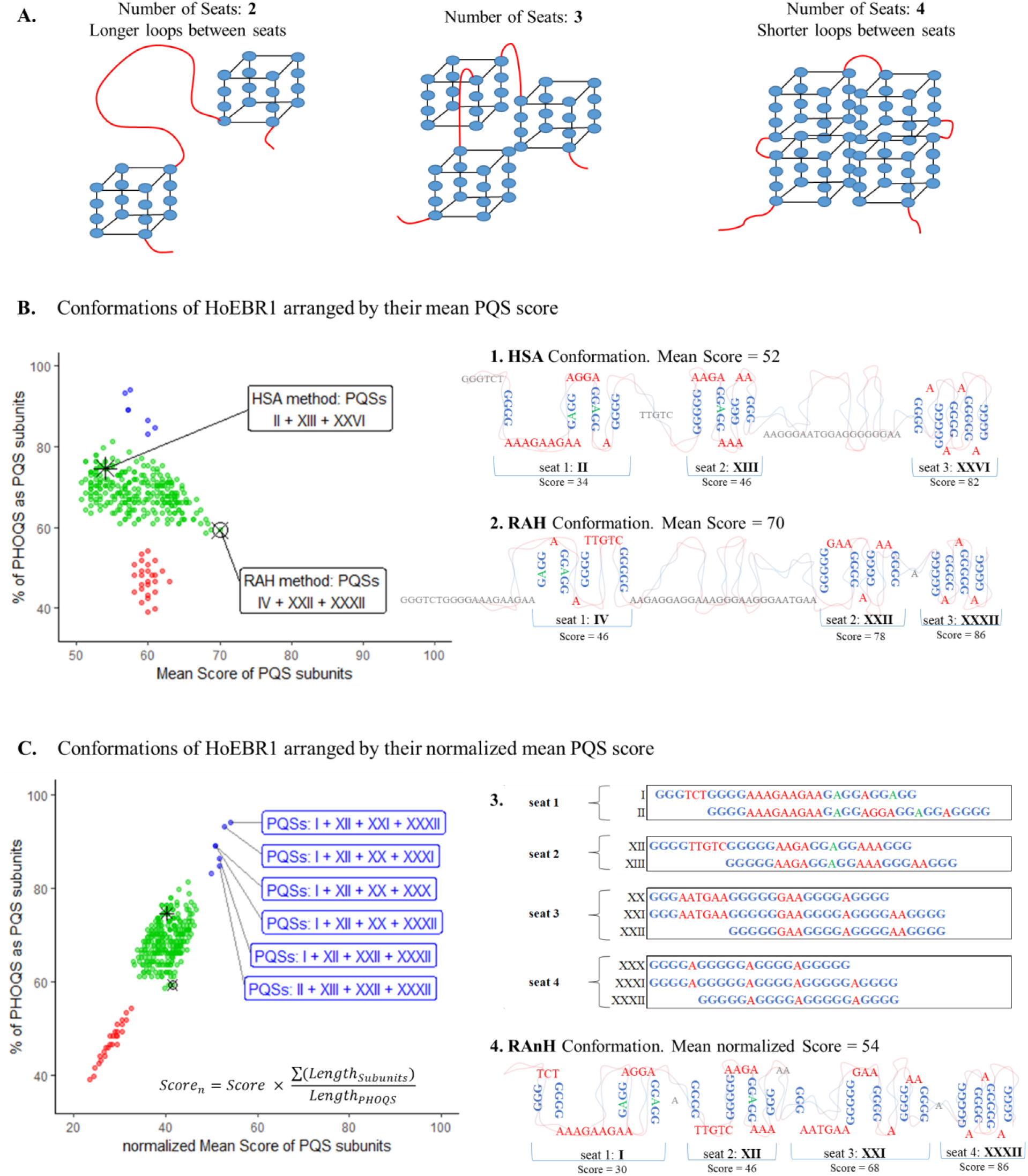
**<u>A.</u>** The PHOQS HoEBR1 can be arranged into up to four *seats*. Greater number of *seats* means smaller loops between units and potentially a gain in structure stability. **<u>B.</u>** 1. High-scoring Sequential Allocation (HSA) conformation is based on assigning *seats* sequentially to the highest scoring candidate and known-to-form G4s. **Graph.** After 10000 iterations of random seat allocation, all 307 candidate conformations of HoEBR1 were found and studied by the mean PQS score of the candidates forming the conformation. In red, blue and green, conformations with 2, 3 and 4 *seats* respectively. **2.** Random Allocation High-scoring (RAH) conformation is the highest mean PQS scoring arrangement. **<u>C.</u> Graph.** The 307 conformations were normalized by the percentage of the PHOQS that is involved as a PQS to favour shorter loops between subunits and greater *seat* density. **3.** The focus was set on the best scoring conformations which present four *seats.* These can be occupied by a combination of 10 candidates. **4.** Random Allocation normalized High-scoring (RAnH) conformation is the mean normalized highest scoring arrangement. Topologies are not accurate.

At first, *seat* allocation was decided to be sequential, assigning a *seat* first to the best scoring PQS with known-to-form G4 in their sequence (method HSA, Highest-score Sequential Assignment). This process yielded a unique organizational candidate that presented three seats and a poor overall score due to election of subunit XXVI as first *seat*. This election ultimately hinders the formation of two other interesting subunits in the tail of HoEBR1 which lowers its overall score (Figure 3, B: 1. HSA Conformation).

An alternative method based on randomly assigning *seats* to candidates was also developed and used. After 10000 iterations, the process identified all possible 307 subunit conformations that can give rise to HoEBR1. This was repeated ten times to make sure no conformations had been excluded. The 307 arrangements were then analysed by their mean *seat* PQS scores (Figure 3, B: Graph), as highest PQS scores are more probable to form G4 *in vitro* and therefore more probable to be the actual PHOQS subunits. Under such pretences, the highest mean score conformation is a three-*seat* structure composed by the PQSs: IV, XXII and XXXII (Figure 3, B: 2. RAH Conformation).

The RAH conformation is based solely on PQS scores and therefore does not consider the loop size between subunits in its study. It can be argued that (as happens within G4s) longer loops are likely to decrease overall stability of the greater structure. Hence, the scores of the conformations were also normalized by the percentage of the PHOQS which is involved as PQS for that given conformation (Figure 3, C: Graph). This way the method discriminates bigger loops between G4 in favour of higher PQS-density conformations. When applied to HoEBR1, 6 four-*seat* configurations scored highly (Scoren > 50, 2 % of total conformations; Figure 3, C), which are the results of electing 10 possible subunits (Figure 3, C: 3). The highest scoring (normalized) arrangement found this way was the combination of the PQS Candidates: I, XII, XXI and XXXII PQS (Figure 3, C: 4. RAnH Conformation). Here, over 96 % of its nucleotides are involved as PQSs and less than 3 % are loops between *seats*.

### Potential G4 relevance in the genome of humans and other organisms

We used G4-iM Grinder’s results to analyse macroscopically the human genome. PQS and PiMS densities (per 100000 nucleotides) were analysed, taking into account both the overall and the most probable to form sequences (those that score at least 40; Figure 4, A). In addition, the genomic quadruplex uniqueness and the number of already confirmed G4s and i-Motif structures detected were calculated. The complete average genomic human density was then used as a reference to compare the human chromosomes. The search was then extended to other species that cause mortal and/or morbid pathologies in humans, including: viruses, fungi, bacteria and parasites. In doing so, we wanted to identify the diseases which could potentially be most effectively treated by targeting these therapeutic structures. We decided to use the genomic sequence density as the means to compare chromosomes and genomes because of the huge length differences between them all. Doing so enabled the most practical and simple method to efficiently and clearly compare the prevalence of these potential structures independently of the genomic length.

**Figure 4.**
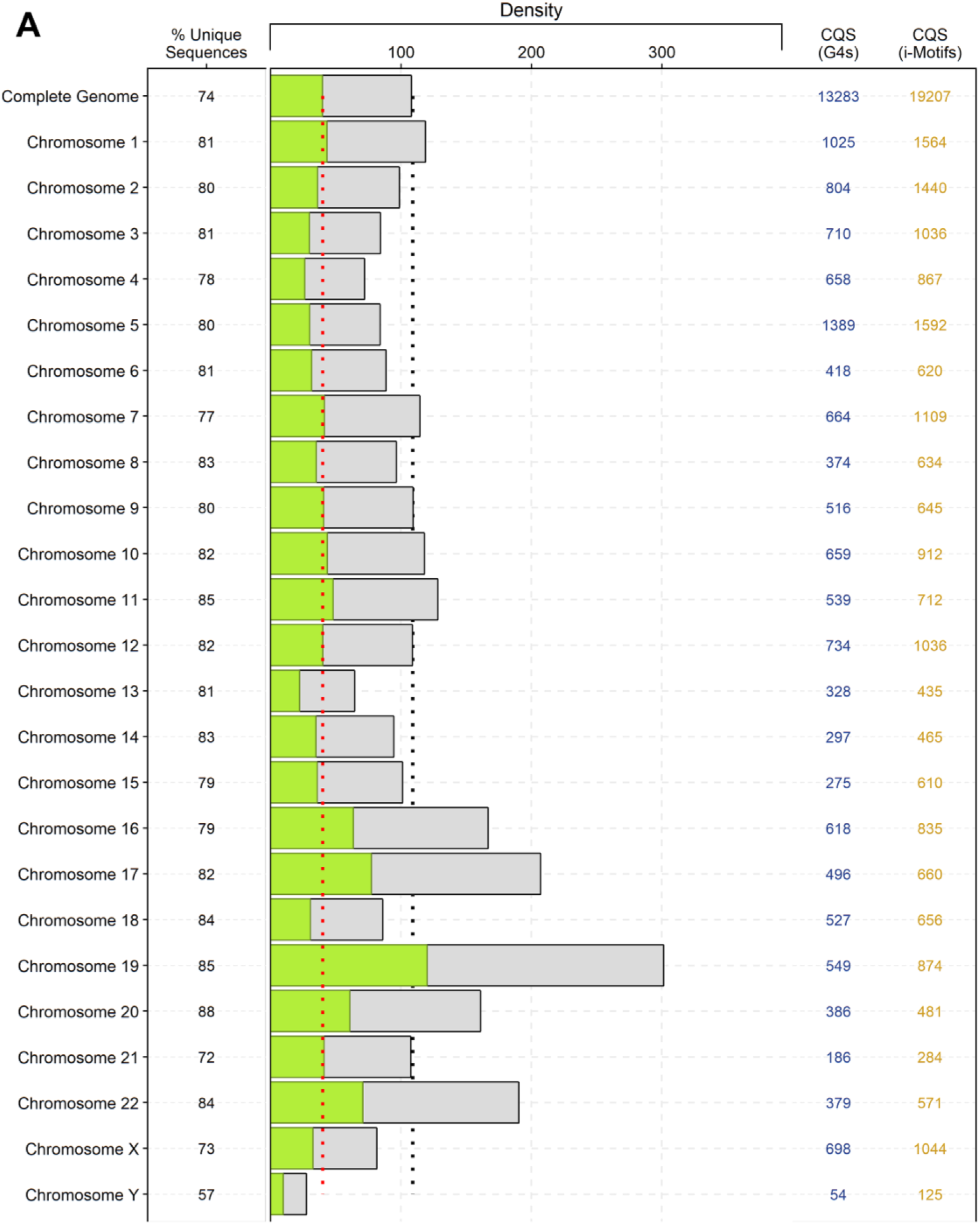
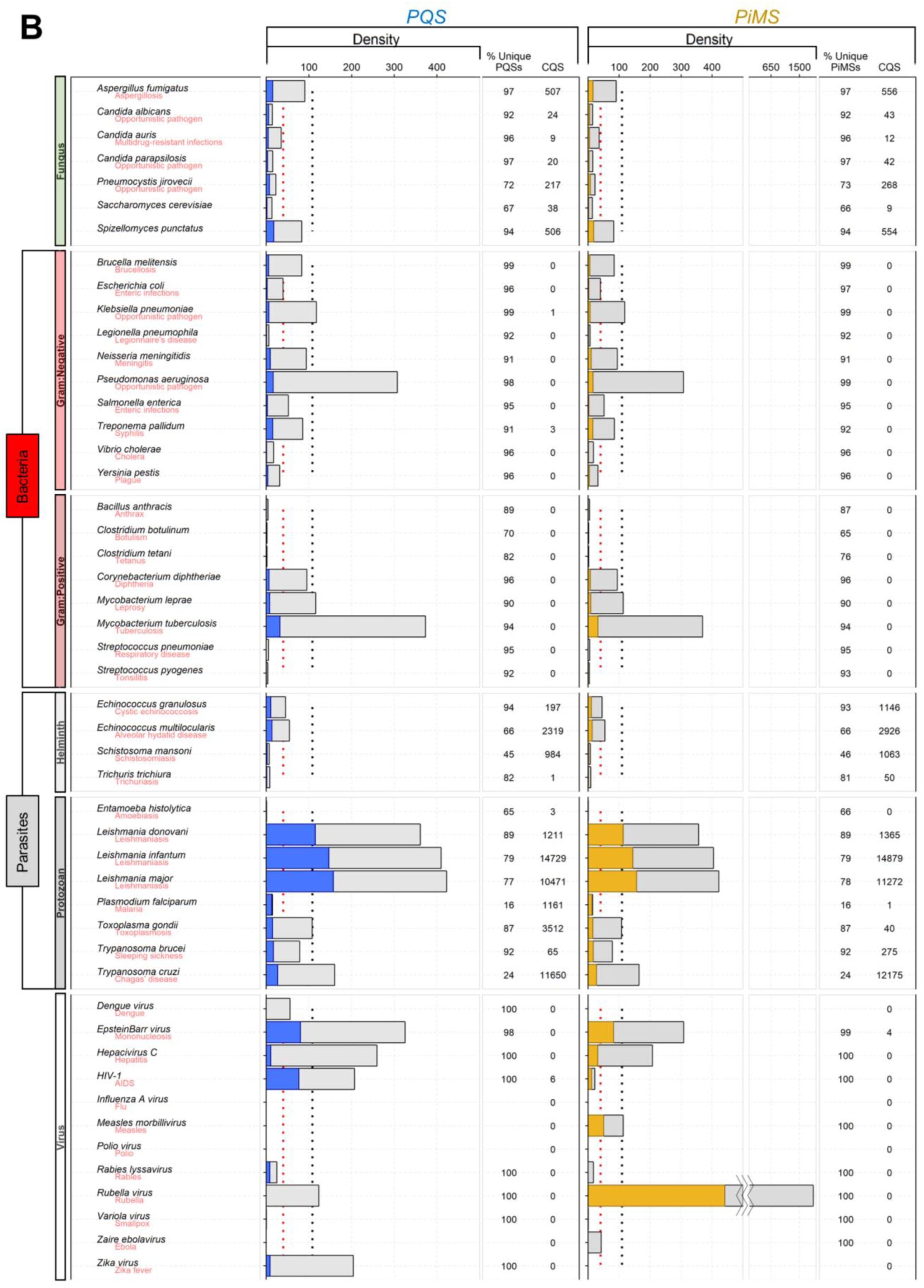
Method 2A (M2A) percentage of unique sequences (sequences which have an occurrence frequency of 1 calculated by the formula: [(M2B results which frequency is 1)/(M2A results)]), PQS and PiMS-densities (per 100000 nucleotides, calculated by the formula: [(M2A results)/(Genome length)] x 100000) and number of known-to-form G4 and i-Motif sequences (abbreviated CQS) detected in humans (**<u>A</u>**) and various other organisms (**<u>B</u>**). Grey bars are the genomic PQS density whilst coloured bars are the genomic density filtered by score of at least 40. Black dotted line is the human average density, and red dotted line is the human average density, which scores at least 40.

The combination of applying different scoring criteria and analytical methods to this -first of its class-quadruplex study allowed a wider context of result interpretation.

Depending on the G4-iM Grinder method employed and the scoring criteria used, the human genome potential structure density (both PQS and PiMS) oscillates between 10 and 300 per 100000 nucleotides. Chromosome 19 showed the highest density (with over 3-fold the human average) followed by chromosomes 17, 22 and 16. Chromosome Y, by the contrary, revealed the smallest genomic density and the lowest percentage of unique sequences. In general, sequences found in the human genome present high frequency of repetition (with just 74 % being unique) and high chance of *in vitro* formation, being a third of the total results over 40 in score. In all chromosomes, hundreds to thousands of already confirmed G4 and i-Motifs were detected. These values surpass most other species examined. However, some exceptions exist (Figure 4, B).

On the one hand, *Leishmania* (and to a less extent the *Trypanosoma* and *Toxoplasma* genus) have very rich quadruplex genomes with many known-to-form G4 and i-Motif sequences within. In *Leishmania major* for example, over 8000 PQS were detected containing the sequence 22Ag (42) with the motif **GGG**TTA. Also, more than 300 PQS containing T30695 and with less frequency T30177 (38), VEGF (42), Scer21 (48), 26gsc (49), Nef8528 (50), IV-1242540 (47), CEB1 (51), A, CC, C and Bc (52), B-raf (53), A3T (49), 96del (41), 27rap (49) and (TG5T)4 (42) were detected. Regarding i-Motifs, cMyb.S (54), cMyc.PY16 (55), KRC6 (43) and cJun (45) (besides the telomeric i-Motif (44)) were also found. In *T. gondii,* over 3000 sequences containing the known-to-form Ara24-1 (48) with the motif **GGG**TTTA, in addition to C, Bc (52), Chla27 (48) and 93del (41), together with the i-Motifs cMyb.S (54), cJun (45), cMyc.C20T (56) and RAD17.2 (57) were also localized. On the other hand, *Plasmodium falciparum* and *Entamoeba histolytica* (causers of malaria and amoebiasis respectively) displayed very low quadruplex densities because of their high genomic AT content (80.6 and 75.2 % respectively). Still, these sequences identified within *P. falciparum* are the least unique of all analysed as most are different variants of its telomeric sequence, PfTel-with the motif **GGG**TTXA (where X can be any nucleotide). The other helminthic parasites and fungi examined present lower densities that those found in the human genome although all had many known to form G4s and i-Motifs within their genomes.

Gram-positive bacteria display very low genomic quadruplex densities all together. The exceptions are the *Mycobacterium* genus-etiological cause of leprosy and tuberculosis- and the Corynebacterium bacteria, which causes diphtheria. These can surpass and even duplicate the human average. In opposition, Gram-negative bacteria have higher densities for those studied here. *Pseudomonas aeruginosa* is the most outstanding genome in this group with a genome 3-fold denser than its human counterpart. *Brucella melitensis* and *Neisseria meningitides* (causers of brucellosis and meningitis, respectively) follow next in density. Several confirmed known-to-form sequences were also found in *Treponema pallidum*, (40) indicating that G4s are already interesting targets against syphilis.

The viruses analysed display a wide range of unique quadruplex densities. Most of them have different PQS and PiMS densities due to being single stranded genomes. For PQS, The Epstein-Barr virus and HIV present higher densities than the human average whilst Zika, Rubella, Rabies and Hepatitis C viruses have similar or slightly lower densities. Within HIV, the known-to-form sequence PRO1 (58) was located. Other viruses including Ebola, Influenza, Measles and Polio viruses were found to be totally void of PQS. When analysing PiMS, Rubella genome was found to be extremely dense, followed by the Epstein-Barr virus which also presented the known cMyb.S (54) i-Motif. Both viruses have greater densities than the human average whilst Measles, Hepatitis C and HIV viruses have similar or slightly lower densities. In other genomes (including the Polio, Influenza and Zika viruses) no PiMS were found.

## DISCUSSION

G4-iM Grinder is a fast, robust and highly adaptable algorithm capable of locating, identifying, qualifying and quantifying quadruplex DNA and RNA structures. These sequences include potential G-quadruplex, i-Motifs and their higher order forms. The adaptation of three scoring systems through machine learning, the structure frequency analysis and the ability to locate already known-to-form G4s and i-Motifs sequences makes G4-iM Grinder’s workflow a practical and easy way to find, filter and select the most interesting quadruplex therapeutic targets in a genome. Furthermore, the modular design and the extensive freedom of variable configuration of G4-iM Grinder gives the user full control of what and how these quadruplex are located and analysed.

Using G4-iM Grinder, we examined the human genome to find new high scoring and frequent quadruplex G and C-based structures as potential new targets. We also identified the longest and most probable higher order sequences to form, some of which have already several known-to-form G4 sequences within. The longest of these structures can involve thousands of nucleotides and hundreds of possible PQS combinations. For example, we analysed HoEBR1 (a recurrent potential higher order quadruplex sequence with good score) and developed the methodology to calculate the best combinations of PQS subunits to form the structure.

A more macroscopic view of the human genome revealed chromosome 19 as the quadruplex densest chromosome and 13, 18 and Y as the least dense ones (with a fall of nearly 66 %). The human genome is still denser and with less unique sequences than most other species examined. However, some parasites and bacteria, such as those in the *Leishmania* and *mycobacterium* genus, present very high densities surpassing by several fold the human average. Other bacteria, like *Pseudomonas aeruginosa*, *Neisseria meningitides* and *Brucella melitensis* are also very rich in potential quadruplex targets, as are the *Trypanosoma* and *Toxoplasma* parasites. In many of these organisms, we identified several sequences that form G4 and i-Motifs *in vitro*. The bacteria and viruses inspected barely presented these known-to-form sequences because these differ from those listed in the sources used to build the known-to-form database. Still the pathological causers of AIDS, hepatitis C, rubella, zika, measles and dengue fever showed very high densities of unique sequences that also exceed the human average. The sum of all these results reflects the great potential G4s have as therapeutic targets against these diseases that currently kill millions worldwide. Other bacteria, parasites and viruses are poorer in or void of quadruplexes and therefore may require less stringent search criteria to find potential targets (for example accepting G or C-runs of length 2).

Future work includes incorporating G4NN and Quadron (when fully developed) as scoring systems. A Shiny application will be developed for G4-iM Grinder and its subsequent result analysis. The quadruplex database will be maintained online to allow external contributions regarding new *in vitro* quadruplex sequences for their identification within G4-iM Grinder results.

## DATA AVAILABITITY

The package and all the results can be found through GitHub (“EfresBR/G4iMGrinder”). Instructions on how to install and use the package can be located in the Supplementary information – 8. G4-iM Grinder package. Results and related information can be located in the Supplementary information – 9. Genomes used and results with all methods.

## SUPPLEMENTARY INFORMATION

Supplementary information are available online.

## FUNDING

This work was supported by The Spanish Ministerio de Economia y Competitividad (C15- 64275-P).

## ACKNOWLEDGEMENTS

The authors thank B. Belmonte, E. Belmonte-Garcia, M. Soto, M. Arévalo, P. Peñalver, S. Heselden, J. L. Mergny and L. Lacroix for their useful insights regarding this topic.

## 1. G4-iM Grinder’s search and analyser algorithm

The algorithm code is written in R and divided into several parts. After a setup is executed, the genomic sequence is analysed and converted into an acceptable format. Then, the complementary strand (if *Complementary* = TRUE) is created. The core of the algorithm is divided in 4 parts, the first two being the search engines (Method 1A and 1B, M1A and M1B respectively), and the other two analysers (Method 2 for overlapping and size-dependent search, and Method 3 for <u>non</u>-overlapping size-<u>in</u>dependent examination, M2A and M3A respectively).

M1A reads and locates all the possible nucleotide runs in the genomic sequence. It starts by identifying non-overlapping perfect runs (with no bulges) and then proceeds by finding imperfect ones in a greedy manner, meaning that it will look for and preferably accept bigger runs than smaller ones. To work, it requires four variables (examples can be found in the supplementary information – S4.Variables, predefined values and examples) which are: *RunComposition*, *BulgeSize*, *MaxRunSize* and *MinRunSize.* M1A output is then passed to M1B, which identifies the relationship between the runs. The method starts by trying to relate runs with their direct neighbours and if unsuccessful, it will expand the search further until *MaxLoopSize* is reached. If this fails, M1B will calculate if reducing the flanking run sizes whilst within the *MinRunSize* acceptable range can solve the problem. This process is dependent on: *MaxLoopSize* and *MinLoopSize*.

M1B data is then analysed by Method 2 (M2A) if the option is selected. This method will join several linked runs to yield the final structures if they comply with the user-defined parameters: *MaxNRuns, MinNRuns*, *MaxPQSSize, MinPQSSize* and *MaxIL* (maximum total number of Bulges accepted per sequence). This is repeated for each linked run so overlapping PQS are all identified together with their position. If desired, *MaxNRuns* can be set to zero to cancel its use in the search and hence accept sequences with more than four runs.

If potential higher-order structures are to be detected, Method 3 (M3A) will select from the M1B data all the runs that have no link with the previous yet are linked with the proceeding ones. These runs will be considered leaders, to which the algorithm will build structures by following the consecutive links. Contrary to M2A, M3A will continue over the *MaxPQSSize* and *MaxNRuns* limits. Hence, it will continue constructing a potential higher order sequence until a non-linked run is added or the end of the sequence is reached. This method depends on the *MinNRuns* and *MinPQSSize* variables.

Once all M2A and M3A results have been identified, G4-iM Grinder will count each structure with the same sequence to calculate its frequency in the genome. These results are stored in a new table called M2B and M3B respectively.

If the computer can and the user allows (by ceding workers through the *NCores* variable), several functions have been given the capacity of applying parallelized computation. This results in a faster analysis.

## 2. Variables, predefined values and examples

**Table 1:** Variables and predefined values for G4-iM Grinder.

| Category | Variable Name | Predefined value | Explanation |
| --- | --- | --- | --- |
| **Function** | *NCores* | 1 | N° of Cores to cede to the function for parallel computation. |
|  | *Verborrea* | TRUE | Allow the function to update the user with its progress. |
| **Sequence** | *Complementary* | TRUE | Analyse the complementary strand of the imputed sequence. |
|  | *DNA* | TRUE | Identifies whether the Sequence is a DNA sequence. If false, it will assume its RNA. Useful for complementary conversion and *KnownQuadruplex* and *KnownNOTQuadruplex*. |
| **For M1A** | *RunComposition* | “G” | Any character can be selected to determine which nucleotide forms the runs. G for G4, C for i-Motifs, tolerates T, U and A or anything else. |
|  | *BulgeSize* | 1 | Number of tolerated non-*RunComposition* nucleotides within the runs, as to allow runs with bulges. |
|  | *MaxRunSize* and *MinRunSize* | 5 -3 | Determines the maximum and minimum numbers of the *RunComposition* nucleotides to be considered a run. |
| **For M1B** | *MaxLoopSize* and *MinLoopSize* | 10 - 0 | Maximum and Minimum number of nucleotides required to assume relationship between runs. |
| **For M2A** | *Method2* | TRUE | To apply method 2. Search for structures with defined size and runs. Will also give frequency counts of each structure detected (M2B). |
|  | *MaxNRuns* and *MinNRuns* | 0 - 4 | Maximum and Minimum number of linked runs which are necessary to form a structure. *MaxNRuns* can be set to 0 to avoid using it as a limiting condition for structure formation (to rely just on structure size). |
|  | *MaxPQSSize* and *MinPQSSize* | 33 - 15 | Maximum and Minimum number of nucleotides which can be part of the final structures. |
|  | *MaxIL* | 3 | Total maximum numbers of bulges allowed for the whole structure. |
| **For M3A** | *Method3* | TRUE | To apply method 3. Search for structures with unrestricted size and numbers of runs. Useful for searching higher forming structures. Depends on variables: *MinNRuns* and *MinPQSSize*. |
| **Quantification** | *LoopSeq* | G, C, T, A | Quantification of each element of *LoopSeq* in all the PQS found (as a % of total length of the structure). Multicharacter values are accepted (like GGG or TTA) |
| **Qualification** | *KnownQuadruplex* | TRUE | Identification of known-to-form-quadruplex (DNA or RNA, G4s or i-Motifs). |
|  | *KnownNOTQuadruplex* | FALSE | Identification of known-NOT-to-form-quadruplex (DNA or RNA, G4s or i-Motifs). |
| **Scoring** | *G4Hunter* | TRUE | To apply G4-hunter evaluation system to the detected final sequences.(1) The results are transformed into a percentage, from 100 to -100 %. |
|  | *PQSfinder* | TRUE | To apply an adaptation of the PQSfinder Scoring system described by Hon et al.(2) It is dependent on *RunComposition* = G or C. |
|  | *cGcC* | FALSE | To apply the cGcC system adopted from Beaudoin *et al.*(3) for RNA sequences. It is dependent on *RunComposition* = G or C. |
|  | *WeightParameters* | 50, 50, 0 | Weighted values of each scoring system as to calculate the final score through weighted averages. Order of vector: G4hunter, PQSfinder & cGcC. |
|  | *FreqWeight* | 0 | Weight of the structure frequency in the final score value for scoring calculations of M2B and M3B. |

### Variable Examples

M1A examples:

| Variables | Sequence | Result |
| --- | --- | --- |
| *RunComposition* = "G" | GGG | Accepted |
| *MinRunSize* = 3 | GG | Non-accepted |
| *BulgeSize* = 0 | GGAG | Non-accepted |
| *RunComposition*: "G" | GGGG | Accepted |
| *MinRunSize*: 4 | GGAG | Non-accepted |
| *BulgeSize*: 1 | GGAGG | Accepted |
| *RunComposition*: "C" | CCC | Accepted |
| *MinRunSize*: 3 | CTTCC | Accepted |
| *BulgeSize*: 2 | CTC | Non-accepted |

M1B examples:

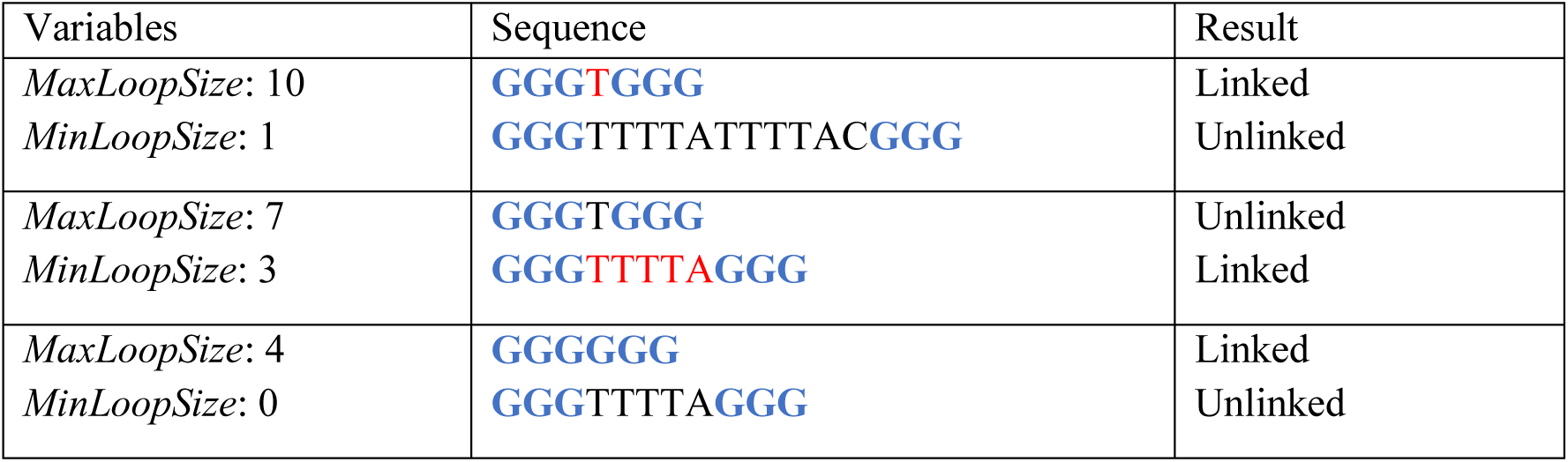

M2A examples:

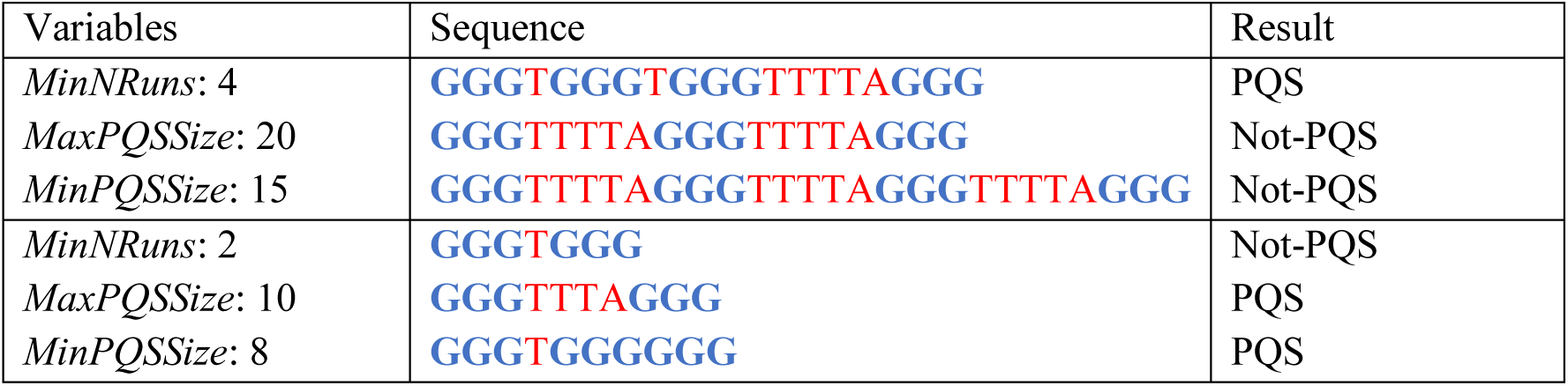

## 3. G-Quadruplex and G4-iM Grinder

392 sequences from the supplementary material of G4Hunter were evaluated to calculate the classification performance of G4-iM Grinder’s Method 2 using predefined parameters. This list of sequences is composed of 94 genomes which do not form G4 and 298 that do.

When the algorithm was applied to the non-forming G4 sequences, 93 of 94 sequences were not recognised as PQS. Only X3CGC with the sequence **GGG**CGC**GGG**CGC**GGG**CGC**GGG** was falsely recognised as a potential quadruplex with fairly high scores given by G4Hunter and PQSfinder (mean score = 48).

When applied to the G4-forming sequences, 233 out of the 298 were correctly recognised with G4-iM Grinder’s predefined values whilst 65 were not. The 5 main reasons why these were not detected are as follows (Table 2):

1. *MinNRuns* < 4: Predefined search values require the presence of at least four G-runs to accept a sequence as a PQS. Therefore, G4s that are intermolecular (with less than 4 G-runs) are not recognised when analysed independently. However, most of these sequences have been found as part of bigger structures.
2. *MinRunSize* < 3: Predefined search values require 3 guanine G-runs to be detected. Hence, neither G-runs of size 2 nor the structures they form were recognised.
3. *BulgeSize* > 1 & *MaxIL* > 3: Predefined search values require sequences to have just 1 bulge per G-run and no more than 3 total bulges per sequence.
4. *MaxLoopSize* > 10: Predefined search values require sequences to have loops of 10 nucleotides or smaller.
5. *MaxPQSSize* > 33: Predefined search values require sequences to have a total length of no more than 33 nucleotides. However this reason can be avoided by analysing the sequences also with the size unrestricted Method 3 (M3).

These search variables can be modified to include these characteristics in any G4-iM Grinder analysis.

This analysis with the predefined variables and its confusion matrix is:

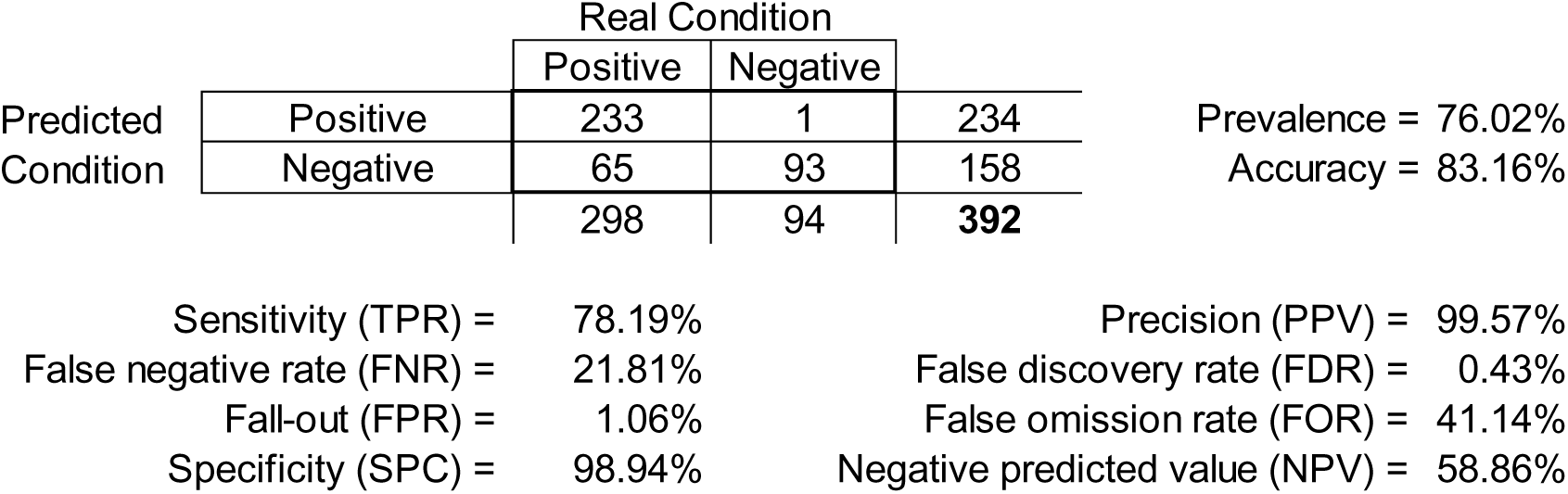

The scores (mean of G4Hunter and G4-iM Grinder’s PQSfinder) of the 233 Sequences were then examined. The mean score of all these structures was of 52.96. Of these structures, 75 % presented a score of 48 or more, and 98 % (228 of 233) scored more than 40. Hence, the filter used in this article to quantify the most probable to form PQS (Score ≥ 40) scored at least the same as 97.9 % of all verified to form sequences examined.

**Table 2:**
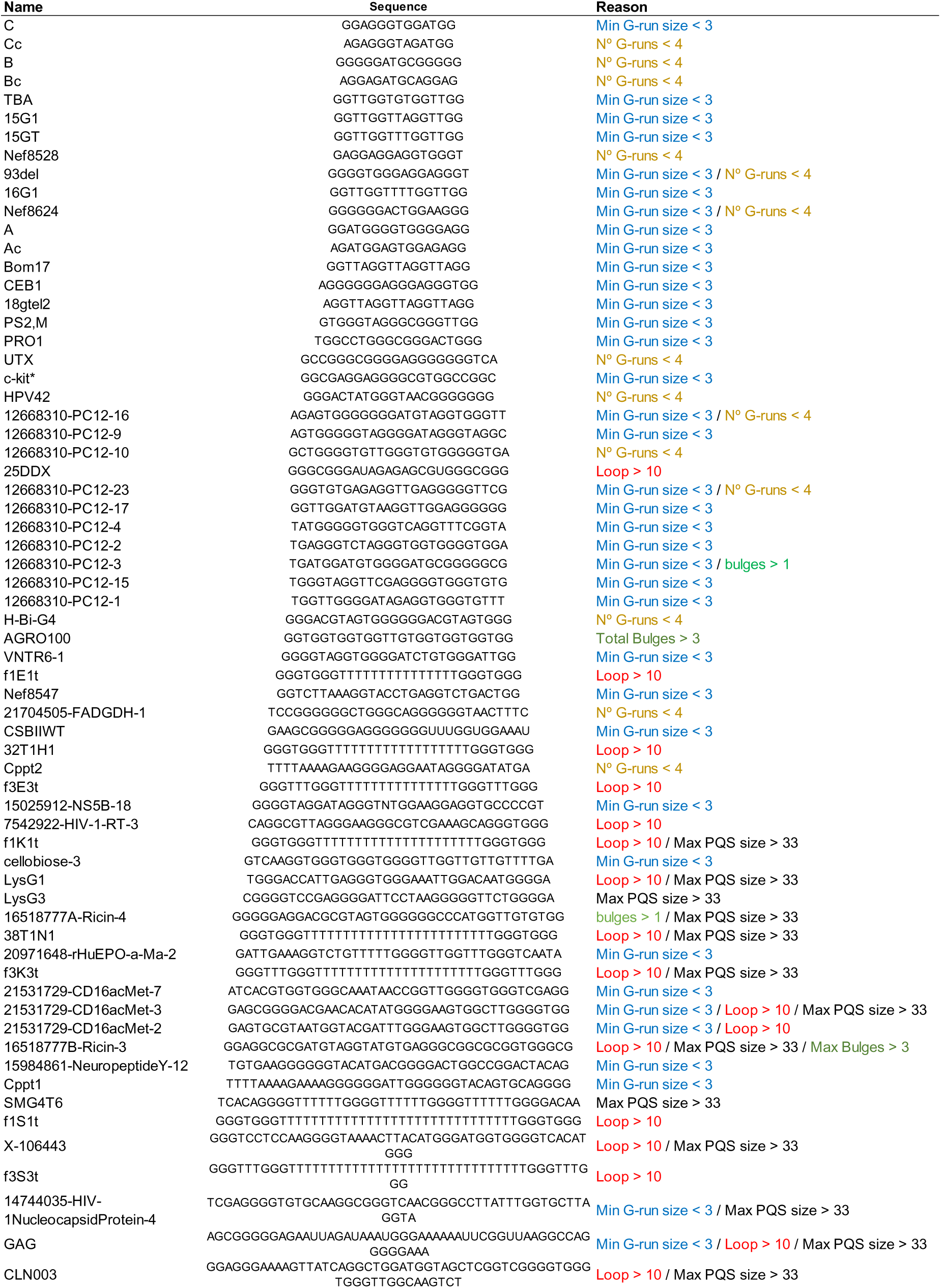
Verified G4 sequences not recognised by G4-iM Grinder using predefined search parameters. The reason why the Sequence was not recognised is given in the third column.

## 4. i-Motif and G4-iM Grinder

100 confirmed i-Motifs were documented from literature and referenced. pH 5, 6, 6.5 and 7 melting temperatures (Tm) were also recorded if stated, although experimental conditions were disregarded. These 100 i-Motifs were analysed by G4-iM Grinder. 77 of the detected sequences were found using G4-iM Grinder’s predefined parameters. 4 were not detected because they present 3 or fewer C-runs (*MinNRuns* < 4) and hence are intermolecular i-Motifs, or are too small (*MinPQSSize* < 15, Table 3). 9 more were detected with the size independent method 3 (M3) as they surpass the max. number of nucleotides (*MaxPQSSize* > 33) for M2. 11 i-Motifs have C-runs of size 2 (*MinRunSize* = 2), and 7 longer loops (*MaxLoopSize* > 10) than accepted. G4-iM Grinder’s i-Motif Sensitivity and False Negative Rate with predefined values is 77 and 23 %. However, all these search variables can be modified to locate all these characteristics in any G4-iM Grinder analysis.

**Table 3:**
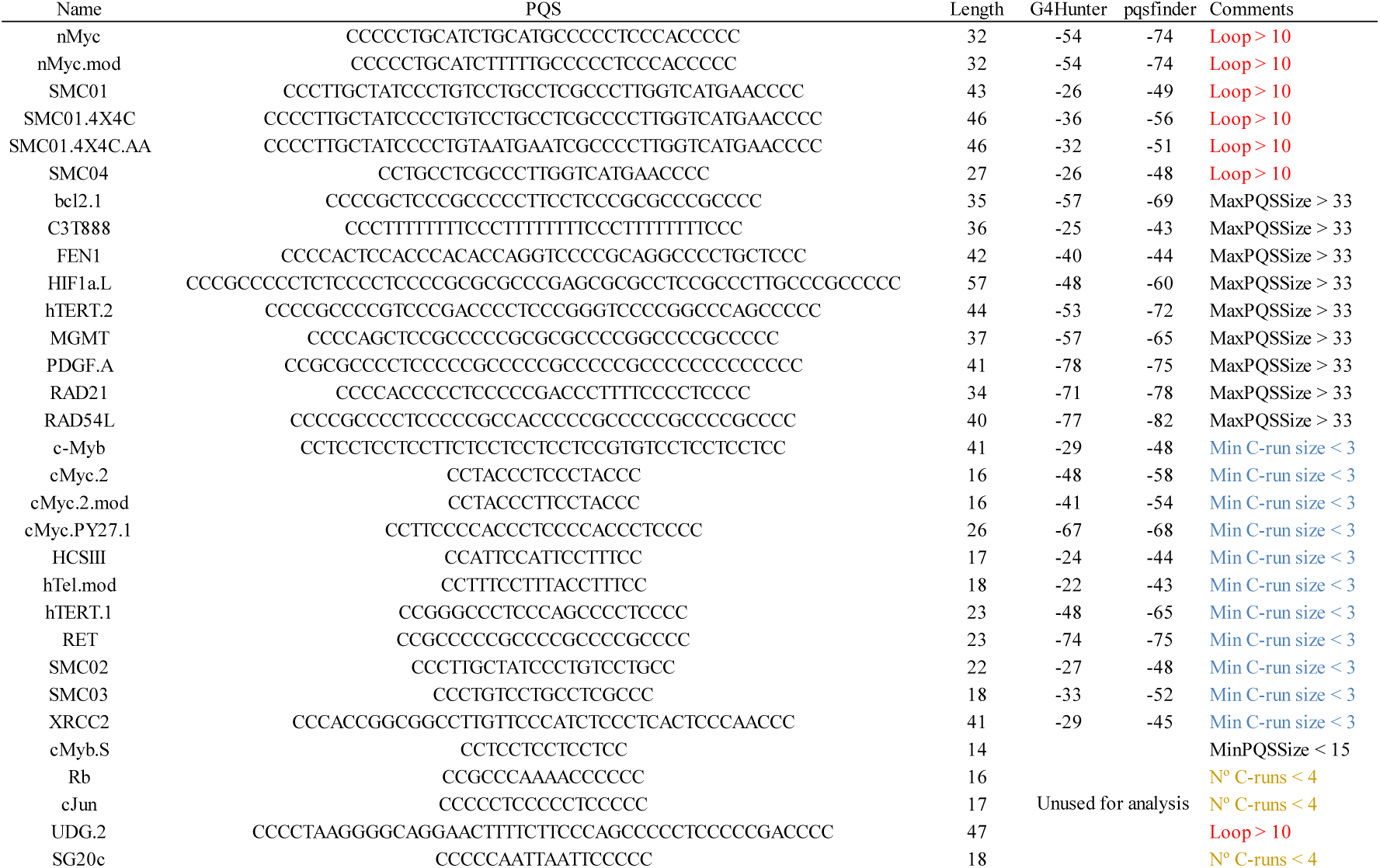
Verified i-Motifs sequences not recognised by G4-iM Grinder using predefined search parameters. The reason why the Sequence was not recognised is stated in the comments column.

## 5. Scoring models and their adaptations

G4-iM Grinder is capable of evaluating potential quadruplex sequences using several published and contrasted scoring algorithms. cGcC was adapted and PQSfinder was upgraded to fit G4-iM Grinder’s workflow. G4hunter was directly implemented from the original article’s supplementary material.

### a) cGcC

cGcC was designed by Beaudoin *et al*. (3) to evaluate RNA PQS by using the formula:

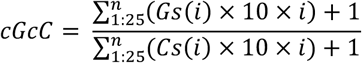

Where Gs(*i*) or Cs(*i*) are the number of G or C elements which compose a run, and *i* is the number of runs. A range of 50 nucleotides are analysed before and after the potential quadruplex sequence and the presence of complementary base nucleotides diminishes the final score. An example is GGGCCCGGG which would score:

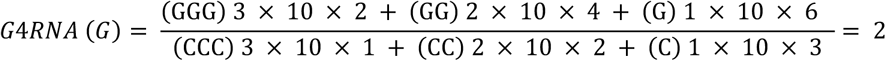

Given the nature of this scoring system, cGcC results are uncontained in the −100 to 100 range and therefore we recommend using cGcC independently of other score systems in the final global scoring evaluation (modulated by the *WeightedParameters* variable).

The >2000 sequences and results found in G4RNA’s supplementary material (4) were compared to G4-iM Grinder’s cGcC function. The results show an almost perfect correlation between both scores (Figure 1).

**Figure 1:**
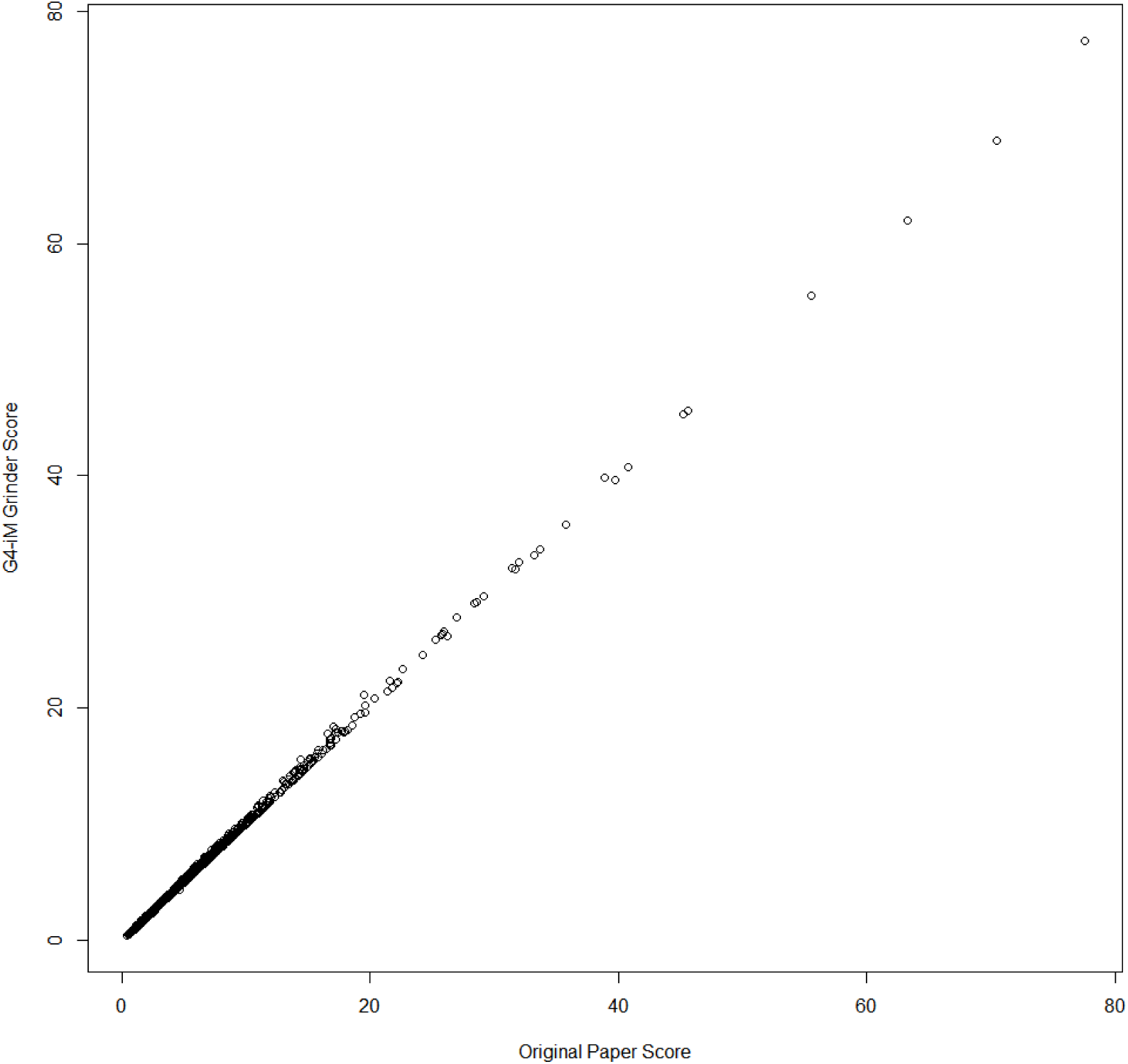
comparison of cGcC results between the G4RNA article and those obtained by G4-iM Grinder cGcC function.

### b) PQSfinder

The original PQSfinder scoring system (2) is based on a modulated formula which analyses quadruplex characteristics such as runs, bulges and loops.

Several modifications were made to adapt it to G4-iM Grinder and to allow greater freedom of analysis. This is because the original function does not allow i-Motif, higher order sequence, or evaluation of sequences with more than four G runs. Low scoring sequences are also not returned.

To adapt the original formula, the appendix of the G run bulges penalisation variables *Fb* (bulge length penalisation factor), *Lbi* (the length of the i-th bulge) and *Eb* (bulge length exponent) were simplified to 1. The number of tetrads (Nt) variable was substituted by the average run size (mNt, excluding from this calculation the bulges) to better analyse longer sequences. Also, the bulge penalisation factor was modulated by multiplying it by the relationship between the minimal and the actual N° of Runs. This was done to avoid scoring distortions in larger sequences due to the characteristic dependency on the absolute number of bulges. Additionally, all segments of the formula were modulated by supplement and exponential constants. The formula used was:

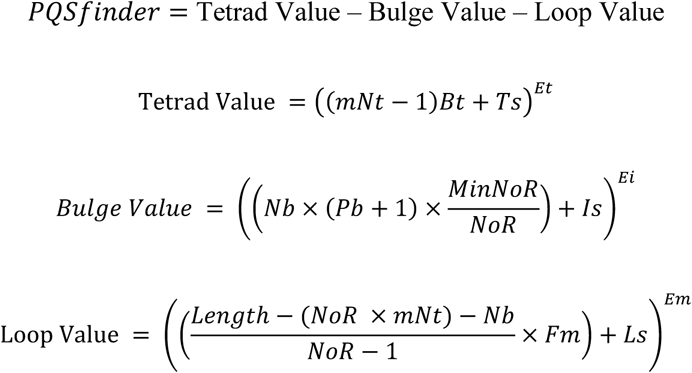

Where *mNt* is the mean run size, *Nb* is the number of bulges within the runs, *Length* is the number of nucleotides in the potential quadruplex and *NoR* the number of runs in the sequence.

G4-iM Grinder’s PQSfinder formula required a re-evaluation to find values that give the nearest score to that of the original PQSfinder, and hence to their *in-silico* G4 formation criteria. The formula was assessed in the non-chromosomal human DNA sequence using both the original PQSfinder and G4-iM Grinder’s PQSfinder whilst accepting only structures with 4 runs (PQSfinder only locates these). 11359 PQS were used to train the parameters (as another 7053 had to be discarded due to the scoring restrictions of the original PQSfinder) by developing and using a machine learning algorithm capable of finding the best values which give the minimal score differences. The range of examination and the best values found are stated in Table 4.

**Table 4:** PQSfinder variables and their values for both the original and G4-iM Grinder’s PQSfinder algorithms. The tested optimization range is displayed in the last column and results of the process are shown underneath.

|  | Name | Description | Original Values | Recommended New Values | Range examined |
| --- | --- | --- | --- | --- | --- |
| Original Constants | *Bt* | Tetrad stacking bonus constant | 40 | 14 | 10 to 50 |
|  | *Pb* | Bulge penalisation constant | 20 | 17 | 10 to 30 |
|  | *Fm* | Loop length penalisation constant | 6.6 | 3 | 0 to 10 |
|  | *Em* | Loop length exponential constant | 0.8 | 1 | 0 to 2 |
| New Constants | *Ts* | Tetrad supplement constant |  | 4 | 0 to 20 |
|  | *Is* | Bulge supplement constant |  | -19 | -20 to 20 |
|  | *Ls* | Loop supplement constant |  | -16 | -20 to 20 |
|  | *Et* | Tetrad exponential constant |  | 1 | 0 to 2 |
|  | *Ei* | Bulge exponential constant |  | 1 | 0 to 2 |
|  | *ET* | Total formula exponential constant |  | 1 | 0 to 2 |
| Results | Mean Difference |  | 4.92 |  |  |
|  | R<sup>2</sup> |  | 0.89 |  |  |
|  | Percentiles of PQS population |  | 50 % - < 3 Scoring difference |  |  |
|  |  |  | 75 % - < 6 Scoring difference |  |  |
|  |  |  | 95 % - < 16 Scoring difference |  |  |

The mean score difference for the same PQS between algorithms was of 4.92 (Original score = G4-iM Grinder score ± 4.92). 50 % of the 11359 PQS analysed fell inside a ± 3 window, 75 % of ± 6 and 95 % of ± 16 (Figure 2). Punctual PQS scoring variations occurred due to different G-run, Loop and bulge definitions, as well as formula variances. However, similarities with the original scoring system are sufficient (R^2^ = 0.89) to allow the effective prediction of quadruplex formation within the original article’s consensus (Figure 3).

**Figure 2:**
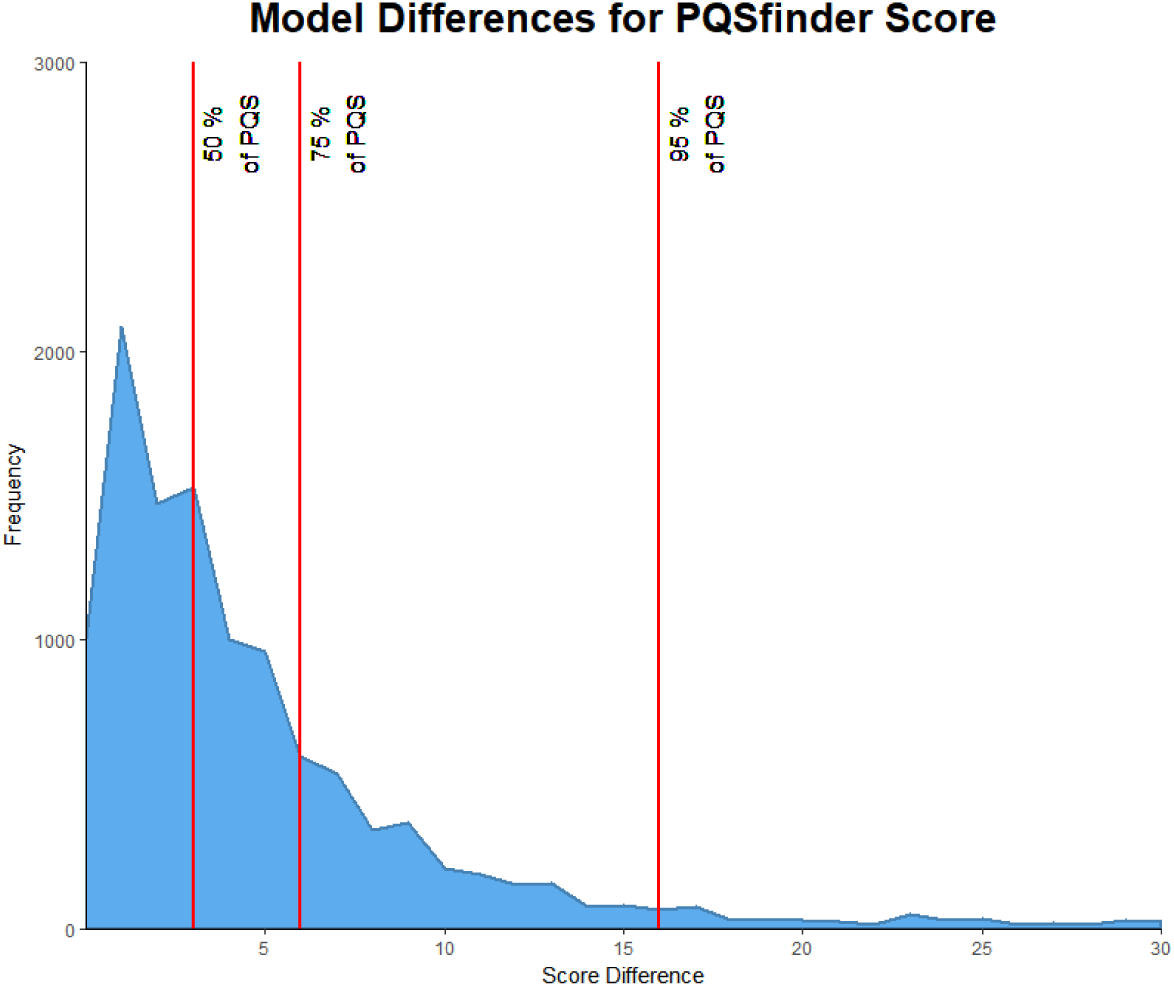
PQS frequencies by their absolute differences between the original and the upgraded G4-iM Grinder’s PQSfinder for 11359 PQS. The horizontal red lines give the accumulative percentage of PQS which difference is inside the range.

**Figure 3:**
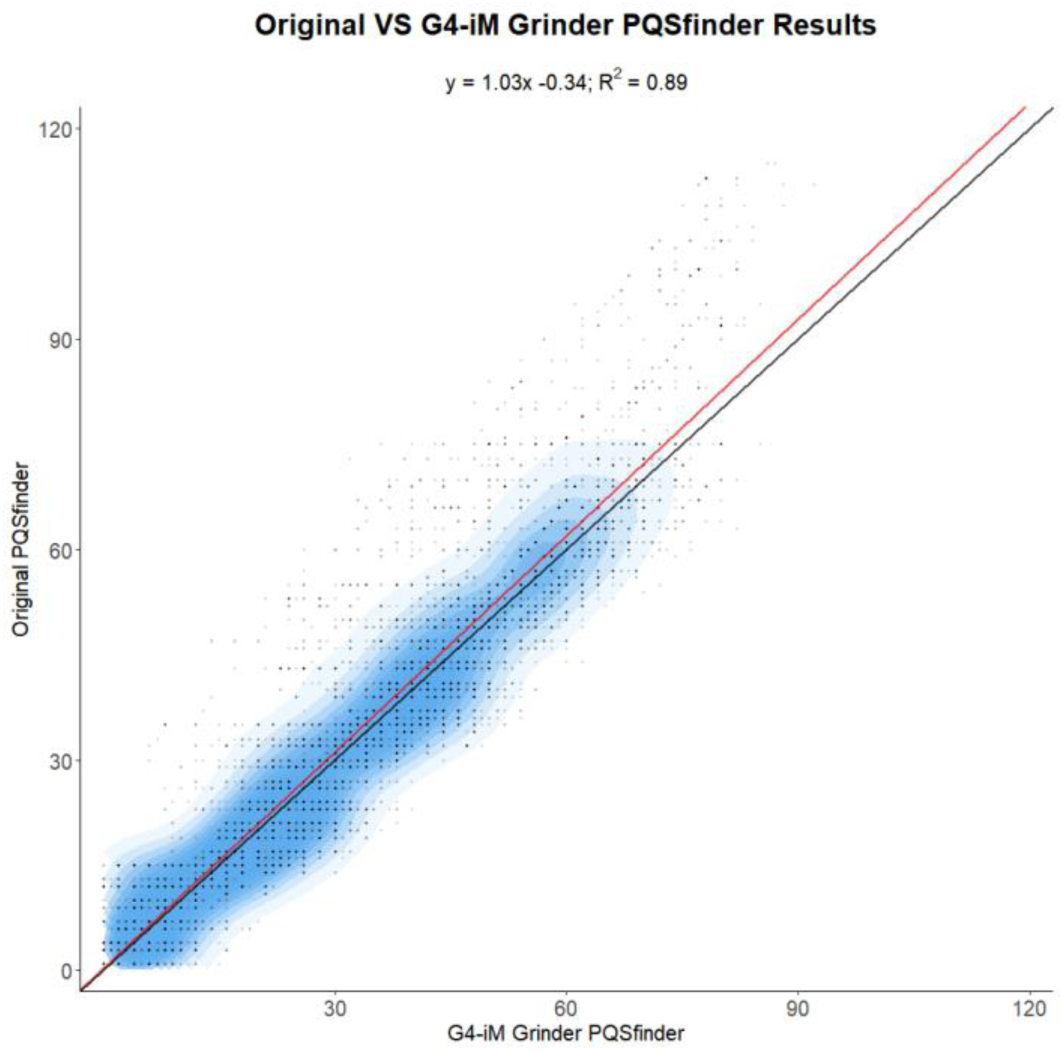
Relationship between the scoring of the original and G4-iM Grinder’s PQSfinder systems for 11359 PQS. The red line is the found correlation line (y = 1.03x - 0.34), and in black the expected perfect match (y = 1x + 0). As a blue shadow, 2D PQS density between both scoring systems.

### c) G4Hunter

This scoring system was directly implemented from the original article (1). However, it was modified to fit into a −100 to 100 % score by multiplying the original score by 25 (the original scale ranges from −4 to 4).

### d) Final Score Evaluation: *WeightedParameters* and *FreqWeight*

After applying all the desired scoring systems, a final score value is calculated modulated by the *WeightedParameters* and *FreqWeight* variables.

*WeightParameters*: is a vector of 3 numbers that state the weight of each scoring system in the final score of a PQS. The first value within the variables is given to the G4Hunter score, the second to PQSfinder and the third to cGcC. Each part is dependent on the activation of its score. This means that if for example *G4Hunter* = FALSE, even if *WeightParameters* = (50, 50, 50), it would calculate the final score assuming the vector is (0, 50, 50).

This value is predefined to be (50, 50, 0) because cGcC has a different scaling system which is not compatible with G4Hunter and PQSfinder. Some cGcC scores can reach values of 2000 and therefore it is recommended to interpret cGcC separately.

*FreqWeigh*t: A constant that gives the importance of the sequence frequency. Useful only for Method 2B and 3B, where frequency of the structures are calculated.

The Final Score is calculated when at least two scoring systems are activated. It will use the results of each scoring system and apply it in a weighted mean formula, where the weight of each value is defined within the variable *WeightParameters*. For Method 2A and 3A, this is:

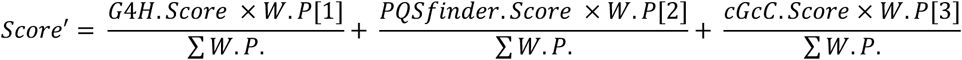

Where *G4H.Score* is the Score of G4Hunter, *W.P*. is the *WeightParameter* variable

For Method2B or Method3B an appendix is added to the formula which takes into account the frequency of the structure using *FreqWeight* to modulate the importance of this characteristic.

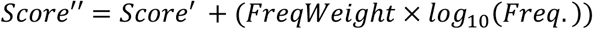

Where *Freq.* is the frequency of the potential quadruplex.

### e) Scoring system for i-Motifs

As G4Hunter punctuates negatively C presence, it was decided to apply an equal but contrary scale for i-Motif likeliness of forming to that of the G based PQSs. This means that when evaluating under the scoring system implemented here, more <u>positive</u> results will mean a higher probability of a PQS being an actual G4; and on the other hand, larger <u>negative</u> values will mean a higher probability of a potential i-Motif (PiMS) being an actual i-Motif. Hence for the analysis of PiMS (*RunComposition* = C), the cGcC formula is elevated and multiplied by −1 and the PQSfinder value is multiplied by −1 as to adapt them to this i-Motif scale.

G4Hunter and G4-iM Grinder’s adaptation of PQSfinder were used to punctuate the *in vitro* probability of formation of 96 known-to-form i-Motif sequences. When applying these score systems to the verified i-Motifs, the results were:

G4Hunter = −49.5 ± 17.0 PQSfinder = −62.1 ± 10.7 Score(mean) = −55.8 ± 13.6

This confirms the direct relationship between high (absolute) probability formation scores and their actual *in vitro* formation (Figure 4, Top: boxplots). These results are very similar to those observed in G4s, which have been used by the original articles (and G4-iM Grinder) to verify the link between high scores and higher potential of quadruplex potential. Moreover, the available melting temperature (Tm) per confirmed sequences was also used to find any correlations between them, the length and the scores (Figure 5). pH 5 Tm and Score(mean) displayed the most interesting correlation (Figure 4, graphs). In all cases, higher scores correlate with higher Tm values, despite the experimental differences for the Tm acquisition, which limits the conclusion extrapolation.

**Figure 4:**
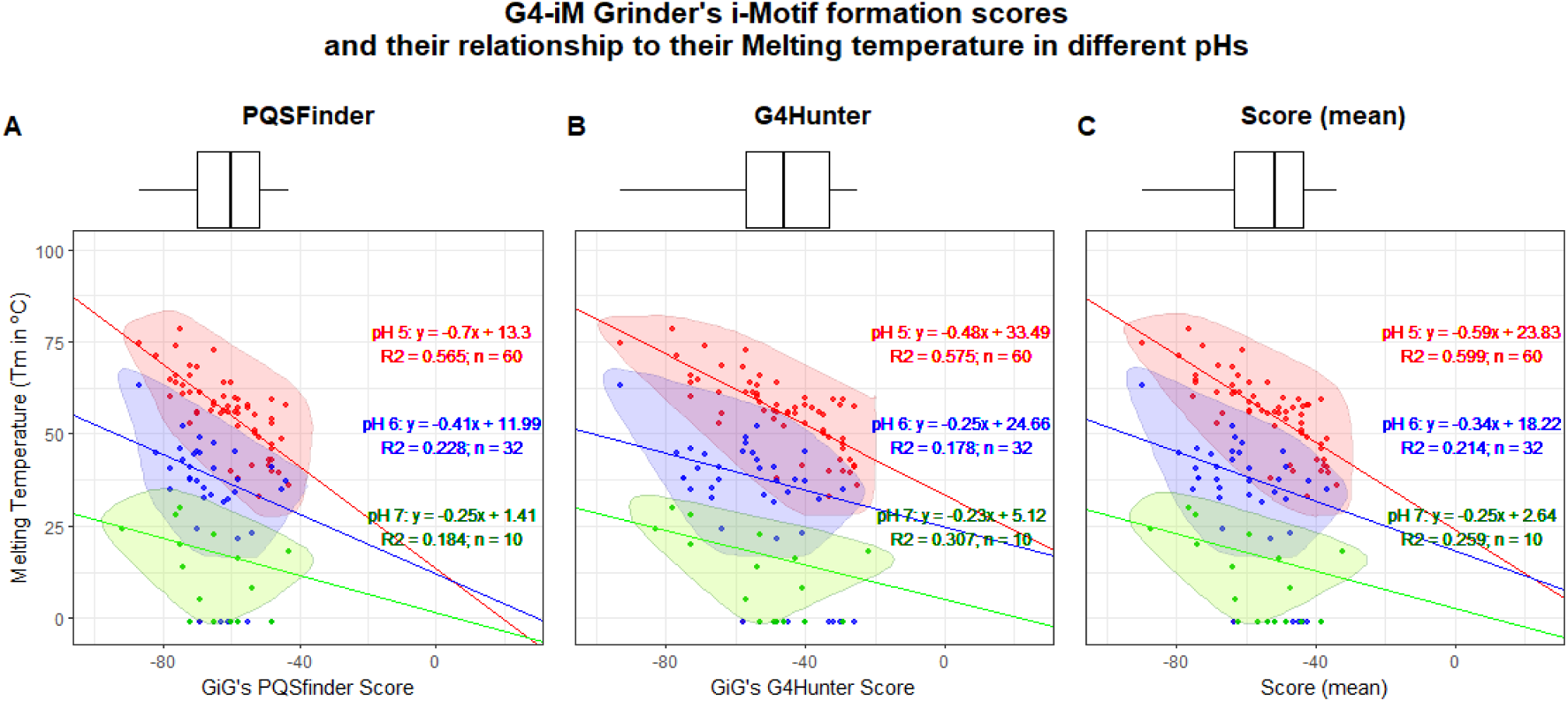
Top: Distribution of i-Motifs regarding their scores. Botton: i-Motif relationship study between their pH-dependent melting temperatures and their PQSfinder score (A), G4Hunter (B) and mean score (C).

**Figure 5:**
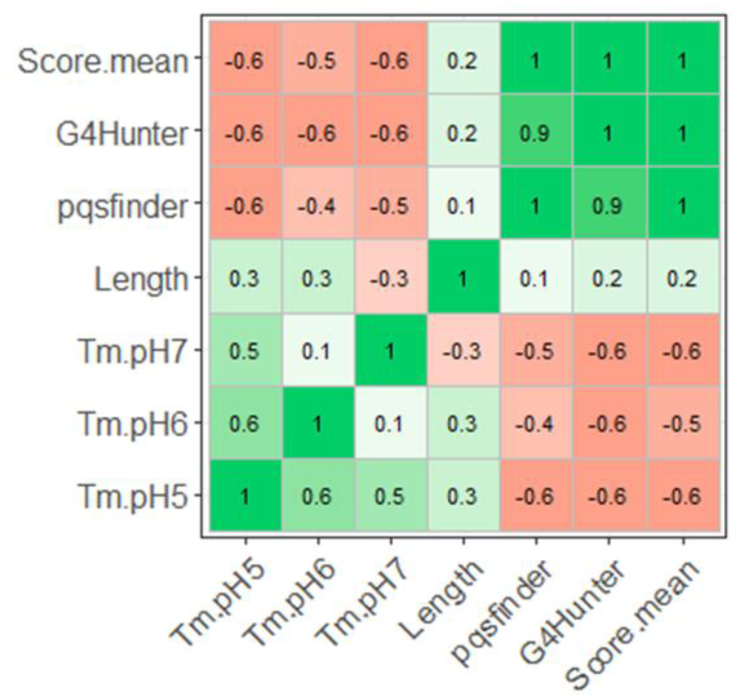
Correlation plot for several characteristics of the 96 i-Motif sequences examined.

## 6. Explanation of other Variables

These variables can be further modified to adapt a G4-iM Grinder analysis.

*Complementary*: If TRUE, it will also create the complementary strand of the sequence and add it to the analysis. A new column will be added to state the sequence’s strand location: original (+) or complementary (-).

*DNA*: Identifies whether the Sequence is a DNA sequence. If *DNA* = FALSE, it means the sequence is an RNA genome. Useful for sequence complementary conversion, *KnownQuadruplex* and *KnownNOTQuadruplex* functions.

These variables will qualify a G4-iM Grinder analysis in a score-independent manner.

*LoopSeq*: A vector which will create a new column per element within the variable to quantify the occurrence of the element in the sequence as a percentage.

For example, if *LoopSeq* = c(G, T, A, C, GGGTTA), it will create 5 columns in the result tables of Method 2 and 3 with the percentage of the final sequence (PQS/ i-motif) which is G, T, A, C and GGGTTA respectively.

*KnownQuadruplex*: A function that will detect *in vitro* confirmed quadruplex within G4-iM Grinder’s Results. For PQS in DNA searches (*RunComposition* = “G”, *DNA* = TRUE) most sequences were downloaded from G4Hunter’s supplementary material (1). For PQS in RNA searches (*RunComposition* = “G”, *DNA* = FALSE), sequences were downloaded from G4RNA DDBB (<u>scottgroup.med.usherbrooke.ca/G4RNA</u>) (4). Several known-to-form DNA i-Motifs (*RunComposition* = “C”, *DNA* = TRUE) were also searched for and included. All these sequences were modified to start and end with the first and last G or C of the sequence as to mimic G4-iM Grinder’s results. Duplicated sequences were eliminated. In total, this DDBB includes 294 DNA and 223 RNA sequences which form G4, and 100 DNA sequences that form i-Motifs. The name/s of the detected quadruplex structures for each sequence are stored in a new column. The number of times detected is stated after the name in between brackets (the search is done in an overlapping manner).

*KnownNOTQuadruplex*: A function that will detect *in vitro* confirmed sequences that do NOT form Quadruplexes within G4-iM Grinder’s Results. For PQS in DNA searches (*RunComposition* = “G”, *DNA* = TRUE) sequences were downloaded from G4Hunter’s supplementary material (1). For PQS in RNA searches (*RunComposition* = “G”, *DNA* = FALSE), sequences were downloaded from G4RNA DDBB (<u>scottgroup.med.usherbrooke.ca/G4RNA</u>) (4). All these sequences were modified to start and end with the first and last G or C of the sequence as to mimic G4-iM Grinder’s results. Duplicated sequences were eliminated as were sequences with less than 18 nucleotides. In total, the DDBB includes 89 DNA and 176 RNA which do NOT form G4. The name/s of the detected non-quadruplex sequences are stored in a new column. The number of times each NOT forming sequences is detected is stated after the name in between brackets (the search is done in an overlapping manner).

## 7. G4-iM-Grinder performance

To test the algorithm, human chromosome 22 (48.5 Mb) was loaded from the ensemble ftp server (version GCA_000001405.25) and analysed with G4-iM Grinder. Performance times and results of all the methods (M2A, M3A and M2B) were analysed with several variable configurations, to measure how these affect code execution (Table 5).

**Table 5:** Results of Chromosome 22 with different parameters. *BulgeSize* = 0, 1 or 2, *MinLoopsize* = 0 or 3, RunComposition = G, T, A or C, and *NCores* = 1 or 7. The rest of the variables were maintained fixed at predefined values, except for cGcC which was changed to TRUE as to evaluate it too and *Complementary* was set to FALSE to measure the differences between the number of structures found within runs of complementary bases. All the analyses were done using an Intel Core i7-4790 K CPU @ 4.00GHz with 16 Gb of RAM.

| <u>Variable configuration</u> |  |  |  |  |  |  |  |  |
| --- | --- | --- | --- | --- | --- | --- | --- | --- |
| *GrunCompositon* | G | G | G | G | G | C | T | A |
| *Ncores* | 1 | 1 | 1 | 1 | 7 | 7 | 7 | 7 |
| *BulgeSize* | 0 | 1 | 1 | 2 | 2 | 2 | 2 | 2 |
| *MinLoopSize* | 0 | 3 | 0 | 0 | 0 | 0 | 0 | 0 |
| <u>Performance (min)</u> |  |  |  |  |  |  |  |  |
| Sequence | 0.01 | 0.01 | 0.01 | 0.01 | 0.01 | 0.01 | 0.01 | 0.01 |
| Method 1A | 0.07 | 0.16 | 0.13 | 0.16 | 0.14 | 0.14 | 0.15 | 0.17 |
| Method 1B | 0.00 | 0.03 | 0.02 | 0.05 | 0.05 | 0.05 | 0.06 | 0.06 |
| Method 2A | 1.35 | 4.47 | 5.73 | 9.91 | 9.15 | 8.80 | 9.98 | 11.66 |
| Method 2B | 0.00 | 0.01 | 0.03 | 0.03 | 0.03 | 0.03 | 0.06 | 0.06 |
| Method 3A | 1.85 | 7.38 | 11.25 | 25.07 | 3.57 | 3.57 | 4.50 | 4.83 |
| Method 3B | 0.00 | 0.01 | 0.03 | 0.03 | 0.03 | 0.03 | 0.06 | 0.06 |
| *G4Hunter* | 0.69 | 1.54 | 3.52 | 4.92 | 0.54 | 0.49 | 0.78 | 0.86 |
| *PQSfinder* | 0.00 | 0.00 | 0.00 | 0.00 | 0.00 | 0.00 | NA | NA |
| cGcC | 0.12 | 0.28 | 0.63 | 0.89 | 0.94 | 0.92 | NA | NA |
| *meanScore* | 0,00 | 0.00 | 0.01 | 0.00 | 0.00 | 0.00 | 0.00 | 0.00 |
| *Quantification* | 0,00 | 0.09 | 0.49 | 0.01 | 0.02 | 0.01 | 0.02 | 0.03 |
| *KnownQuadruplex* | 0.08 | 0.22 | 0.47 | 0.69 | 0.73 | NA | NA | NA |
| **Total** | **4.17** | **14.20** | **22.32** | **41.77** | **15.21** | **14.05** | **15.61** | **17.74** |
| <u>Results - Structures found</u> |  |  |  |  |  |  |  |  |
| Method 2A | 17320 | 35041 | 97098 | 116290 | 116290 | 117428 | 210556 | 228603 |
| Method 2B | 15906 | 32216 | 87690 | 107945 | 107945 | 108154 | 175696 | 183499 |
| Method 3A | 8260 | 24760 | 39421 | 70858 | 70858 | 69714 | 96983 | 103060 |

Both performance time and results show dependency on the sequence analysed, its composition and organization, and the parameters employed. The simplest of options examined here (*BulgeSize* = *MinLoopsize* = 0) yielded 17320 PQS. This is 1.14 million PQS when extrapolated to the whole genome, a number similar to previous estimates for folding rule abiding-structures (5). These results increased fivefold when the number of acceptable bulges (*BulgeSize)* was increased to 1. When they were set to 2, smaller differences were observed because of the user-defined variable *MaxIL* that limits the maximum number of total acceptable bulges in the sequences.

These results were compared to those of other nucleotide-composition runs, including potential i-Motif sequences which are similar in number to G-based PQS. For structures composed of T and A runs, which have no known physical meaning, this count almost doubles that of G and C results. This is in accordance with a previous report by Huppert (5).

Regarding performance, the most time-consuming processes were M2A, M3A and G4Hunter scoring system. The total process time was optimized by increasing the numbers of cores of the computer to do the calculations. Using 7 cores reduced the time by 66 % in total. As a comparison, PQSfinder was downloaded directly from CRAN (December 2017) and executed with the same genomic sequence to compare the execution times. In total, the analysis with PQSfinder took 4 hours and 16 minutes to finish when running with predefined variables (except strand which was set to “+”).

## 8. Grinder Package

G4-iM Grinder can be download from github: EfresBR/G4iMGrinder. G4-iM Grinder requires the installation of other CRAN based and Bioconductor packages. Please, ensure all required packages are installed. G4-iM Grinder was successfully downloaded and tested in MacOS 10.12.6, Windows 10 (x64) and Ubuntu 18.04.1 (x64) running R 3.6.0 (and older) and R studio 1.2.1135 (and older). In Ubuntu, the installation of devtools may require further effort (<u>link)</u>. Other OS including x86 systems have not been tested.

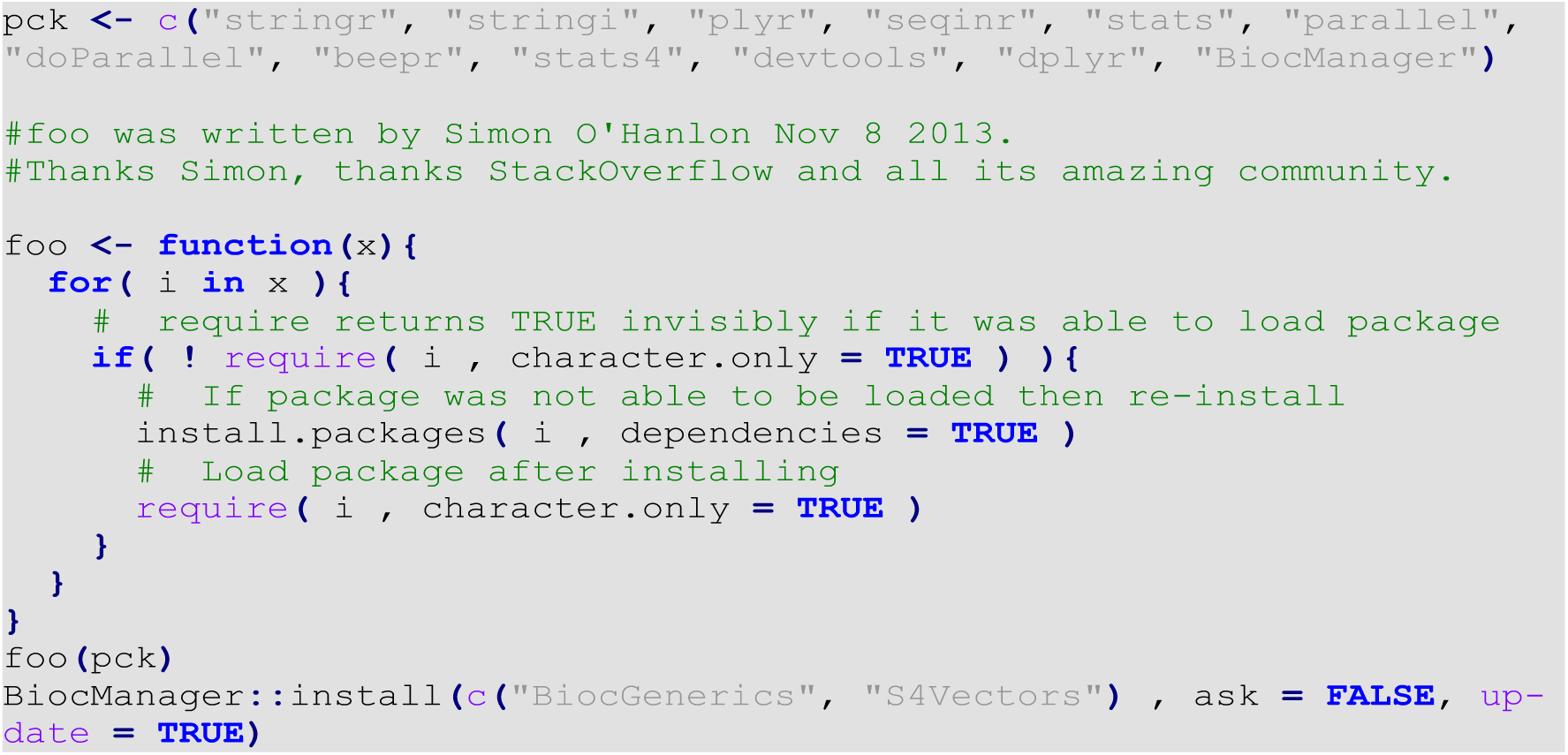

To install and load G4-iM Grinder

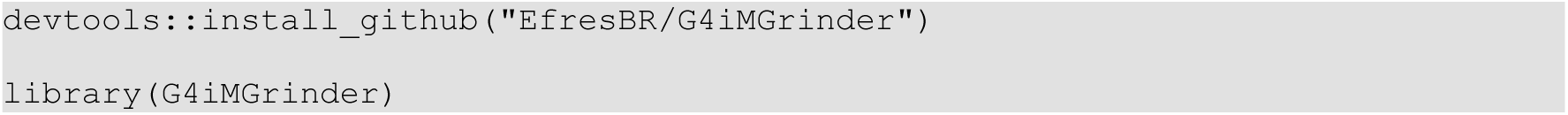

Executing a genomic G-Quadruplex analysis with **G4iMGrinder** function

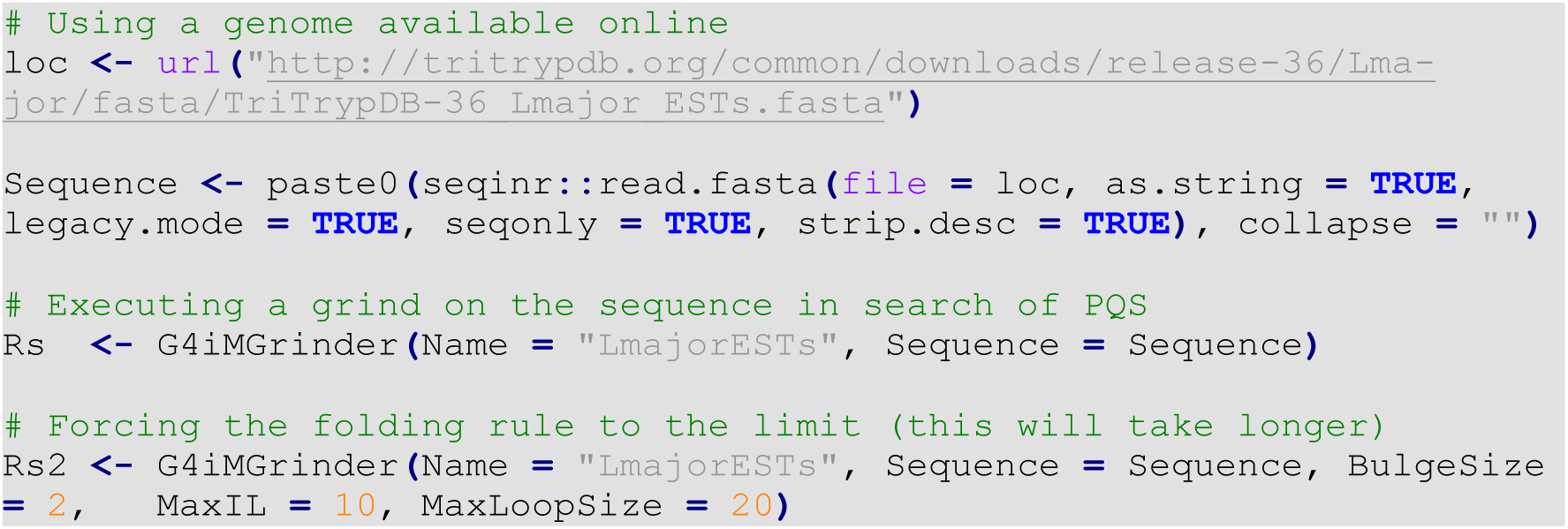

Summarizing an analysis with **GiGList.Analysis** function to compare the results between genomes. This will quantify the number of results and density of each analysis. It will also give the number of results that have at least a minimum frequency, score and size. These variables can be modified. See the package documentation for more information regarding **GiGList.Analysis**.

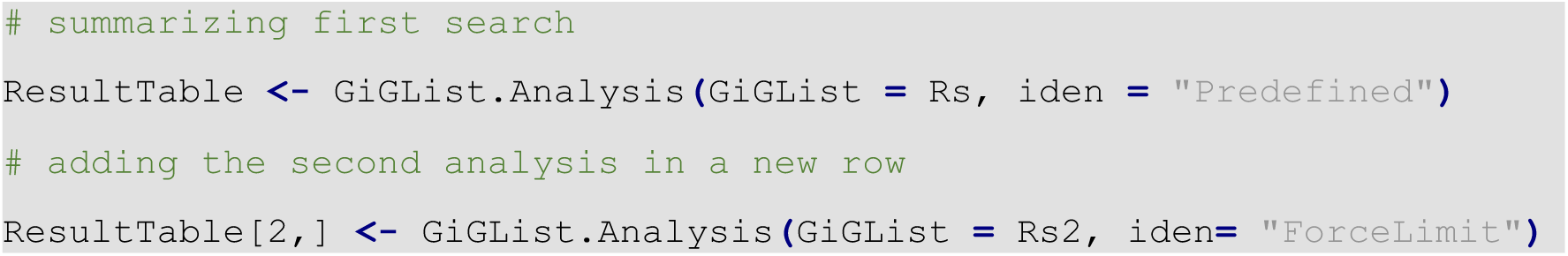

Executing an analysis of a higher order structure with **GiG.M3Structure** to analyse its potential subunit configuration. This will give all and the most interesting subunit conformations as stated in the article. See the package documentation for more information regarding **GiG.M3Structure**.

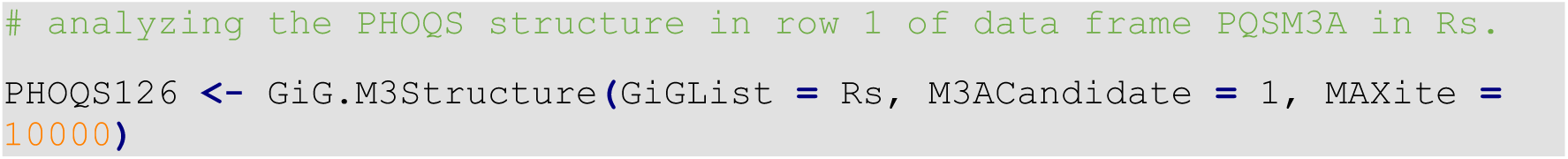

Finding the reference for the Known-To-Form Quadruplex structures of an interesting Result. This procedure is the same for Known-NOT-To-Form sequences.

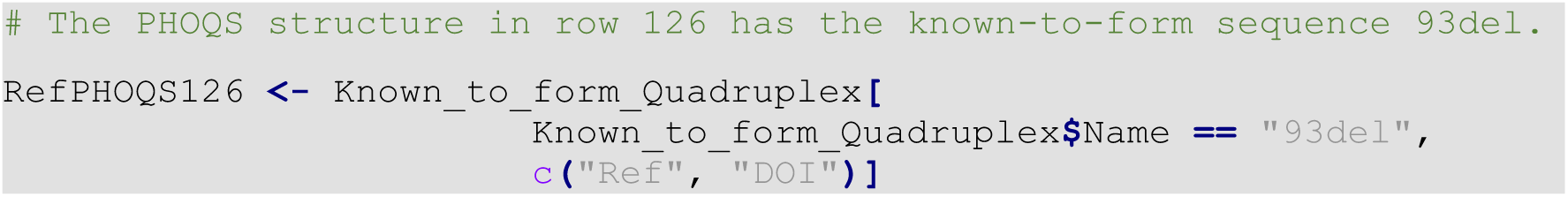

Updating a G4-iM Grinder analysis with different variables using the **GiGList.Updater** function. This will avoid doing a new search analysis on the sequence and therefore will be more time and resource efficient.

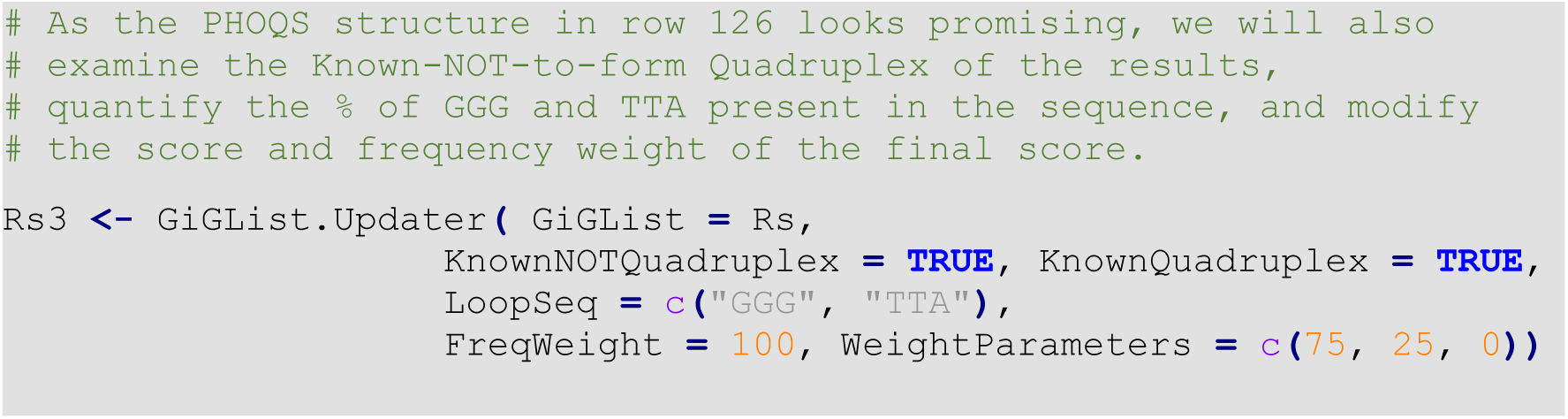

To search for potential i-Motifs in the genome we can repeat the analysis with **G4iMGrinder** function changing *RunComposition* = “C”. However, if a previous analysis of the genome has already been done with the complementary base-pair, we can also use the function **GiGList.Updater** to search for the resulting opposite structures.

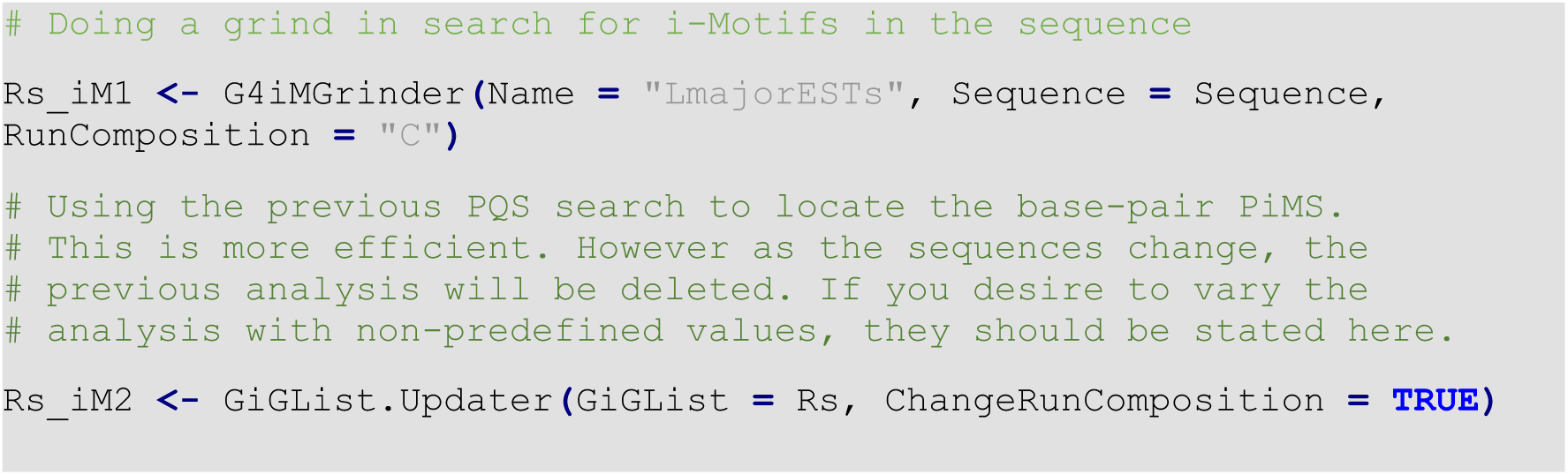

## 9. Genomes used and results with all methods

The complete human chromosomic genome (GRCh38.p12, Genome Reference Consortium Human Build 38, INSDC Assembly GCA_000001405.27– gh38) was downloaded from the Sanger Institute (www.sanger.ac.uk) in May 2019. Other non-human genomes used in this paper were downloaded in May 2019 from https://www.ncbi.nlm.nih.gov/, and include:

| **<u>Name</u>** | **<u>iden</u>** | **<u>RefSeq</u>** |
| --- | --- | --- |
| Aspergillus fumigatus | Fungus | NC_007194.1 to NC_007201.1 |
| Candida albicans | Fungus | NC_032089.1 to NC_032096.1 |
| Candida auris | Fungus | PGLS000000000.1 |
| Candida parapsilosis | Fungus | HE605209.1 |
| Pneumocystis jirovecii | Fungus | LFWA000000000.1 |
| Saccharomyces cerevisiae | Fungus | NC_001133.9 to NC_001224.1 |
| Spizellomyces punctatus | Fungus | ACOE000000000.1 |
| Brucella melitensis | Gram:Negative | NC_003317.1 to NC_003318.1 |
| Escherichia coli | Gram:Negative | NC_011750.1 |
| Klebsiella pneumoniae | Gram:Negative | NC_016845.1 |
| Legionella pneumophila | Gram:Negative | NC_002942.5 |
| Neisseria meningitidis | Gram:Negative | NC_003112.2 |
| Pseudomonas aeruginosa | Gram:Negative | NC_017548.1 |
| Salmonella enterica | Gram:Negative | NC_003197.2 |
| Treponema pallidum | Gram:Negative | NZ_CP003679.1 |
| Vibrio cholerae | Gram:Negative | NC_002505.1 to NC_002506.1 |
| Yersinia pestis | Gram:Negative | NC_003143.1 |
| Bacillus anthracis | Gram:Positive | NC_003997.3 |
| Clostridium botulinum | Gram:Positive | NC_009698.1 |
| Clostridium tetani | Gram:Positive | NC_004557.1 |
| Corynebacterium diphtheriae | Gram:Positive | NZ_LN831026.1 |
| Mycobacterium leprae | Gram:Positive | NC_002677.1 |
| Mycobacterium tuberculosis | Gram:Positive | NC_000962.3 |
| Streptococcus pneumoniae | Gram:Positive | NC_003098.1 |
| Streptococcus pyogenes | Gram:Positive | NC_002737.2 |
| Echinococcus granulosus | Helminth | APAU000000000.2 |
| Echinococcus multilocularis | Helminth | CBLO000000000.2 |
| Schistosoma mansoni | Helminth | NC_031495.1 to NC_002545.1 |
| Trichuris trichiura | Helminth | CBXK000000000.1 |
| Entamoeba histolytica | Protozoan | AAFB000000000.2 |
| Leishmania donovani | Protozoan | NC_018228.1 to NC_018263.1 |
| Leishmania infantum | Protozoan | NC_009386.2 to NC_009420.2 |
| Leishmania major | Protozoan | NC_001905.3 to NC_007287.2 |
| Plasmodium falciparum | Protozoan | NC_004325.2 to NC_037283.1 |
| Toxoplasma gondii | Protozoan | NC_031467.1 to NC_001799.1 |
| Trypanosoma brucei | Protozoan | NC_026734.1 to NC_026744.1 |
| Trypanosoma cruzi | Protozoan | NZ_AAHK00000000.1 |
| Dengue virus | Virus | NC_001474.2 |
| EpsteinBarr virus | Virus | LS992244.1 |
| Hepacivirus C | Virus | NC_004102.1 |
| HIV-1 | Virus | NC_001802.1 |
| Influenza A virus | Virus | NC_007366.1 |
| Measles morbillivirus | Virus | NC_001498.1 |
| Polio virus | Virus | NC_002058.3 |
| Rabies lyssavirus | Virus | KY982923 |
| Rubella virus | Virus | NC_001545.2 |
| Variola virus | Virus | NC_001611.1 |
| Zaire ebolavirus | Virus | NC_002549.1 |
| Zika virus | Virus | NC_012532.1 |

The complete G4iM-Grinder Results used for this article can be downloaded from github: EfresBR/G4iMGrinder (https://github.com/EfresBR/G4iMGrinder). These results are compressed in the archive named “Results.RAR”. Inside, four RData images are included: “*hg39PQS.RData*” (Human G-based PQS), “*hg39PiMS.RData*” (Human C-based PiMS), “NonHumanPQS.RData” (non-human G-based PQS) and “NonHumanPiMS.RData” (non-human C-based PiMS).

For human results, 25 G4-iM Grinder Lists (GiG-Lists) and 1 data frame exist within each Rdata archive. 24 of these lists are the results for each individual human chromosome and the last (and biggest) is the complete and joint M2B and M3B evaluation of the genome (it is **NOT** the sum of the individual chromosomes). The data frame *ResultTable* is a summary of the results, after using G4-iM Grinder’s Analysis function which helps extract the basic information of the results in a fast and convenient way.

For non-human results, each GiG-List is the analysis of an organisms. The data frame *A.ResultsTable* is a summary of the results, after using G4-iM Grinder’s Analysis function. *A.GenomeTable* contains the information of each genome examined.

## Notes

https://github.com/EfresBR/G4iMGrinder

